# Mitotic events depend on regulation of PLK-1 levels by the mitochondrial protein SPD-3

**DOI:** 10.1101/2023.01.11.523633

**Authors:** Yu-Zen Chen, Vitaly Zimyanin, Stefanie Redemann

## Abstract

In metazoans, Polo Kinase (Plk1) controls several mitotic events including nuclear envelope breakdown, centrosome maturation and kinetochore assembly. Here we show that mitotic events regulated by Polo Like Kinase (PLK-1) in early *C. elegans* embryos depend on the mitochondrial-localized protein SPD-3. *spd-3* mutant one-cell embryos contain abnormally positioned mitotic chromosomes and prematurely and asymmetrically disassemble the nuclear lamina. Nuclear envelope breakdown (NEBD) in *C. elegans* requires direct dephosphorylation of lamin by PLK-1. In *spd-3* mutants PLK-1 levels are ~6X higher in comparison to control embryos and PLK-1::GFP was highly accumulated at centrosomes, the nuclear envelope, nucleoplasm, and chromosomes prior to NEBD. Partial depletion of *plk-1* in *spd-3* mutant embryos rescued mitotic chromosome and spindle positioning defects indicating that these phenotypes result from higher PLK-1 levels and thus activity. Our data suggests that the mitochondrial SPD-3 protein controls NEBD and chromosome positioning by regulating the endogenous levels of PLK-1 during early embryogenesis in *C. elegans*. This finding suggests a novel link between mitochondria and mitotic events by controlling the amount of a key mitotic regulator, PLK-1 and thus may have further implications in the context of cancers or age-related diseases and infertility as it provides a novel link between mitochondria and mitosis.

## Introduction

Mitochondria are organelles in which cellular respiration occurs, as well as a variety of other biosynthetic processes, such as the regulation of Ca^2+^, apoptosis, and the production of reactive oxygen species (Jeong and Seol 2008; Tilokani et al. 2018; Amorim et al. 2022). Artificially disrupting mitochondrial function in oocytes and early embryos was shown to cause defects in maturation (Al-Zubaidi et al. 2021), fertilization (Pasquariello et al. 2019), errors in cell division and chromosome segregation (Wang et al. 2020; Mikwar et al. 2020), and loss of viability and defects in metabolism (Al-Zubaidi et al. 2021). These observations suggest a link between mitochondrial defects, spindle assembly abnormalities, chromosome segregation errors and aneuploidy. As mitochondria are strongly involved in metabolism and the main generators of energy in form of ATP it is possible that changes in the available energy can negatively affect spindles. In particular, the function of motor proteins, which are required for spindle assembly and chromosome segregation depends on available ATP (Sweeney and Holzbaur 2018), and changes in the ATP content could impair motor function, ultimately leading to mitotic errors (Cheng et al. 2021). However, the mechanisms of how defects in mitochondria affect spindle function in mitosis have remained elusive.

The *C. elegans* Spindle-defect-3 (SPD-3) protein is a mitochondria localizing protein that was isolated in an EMS-mutagenesis screen in search of cell division mutants (O’Connell et al. 1998). Previous publications have shown that defects in the SPD-3 protein in the nematode *C. elegans* impair homolog pairing by causing a severe reduction in the mobility of SUN-1 aggregates at the start of meiotic prophase (Labrador et al. 2013). It was suggested that this defect is due to the reduced function of cytoskeletal motors in *spd-3* mutants, caused by a defect in mitochondrial function and associated changes in ATP levels (Labrador et al. 2013). This was further supported by the observation of mitotic spindle positioning defects in one-cell *C. elegans spd-3* mutant embryos, which are comparable to spindle positioning defects after dynein depletion (Yoder and Han 2001; Dinkelmann et al. 2007).

Here we show that mutation of spd-3 does not only impair spindle positioning as previously reported but also chromosome positioning and the nuclear integrity during mitosis. Interestingly, we found that Polo-like kinase (PLK-1) levels are significantly increased in the *spd-3* mutant. PLK-1 is a crucial mitotic kinase that functions in many aspects of cell division, such as mitotic entry (Thomas et al. 2016), centrosome maturation (Fu et al. 2015), nuclear envelope (NE) disassembly (Martino et al. 2017; Linder et al. 2017; Dawson and Wente 2017), and asymmetric cell division (Han et al. 2018) in *C. elegans* and mammalian cells.

PLK1 function is not limited to mitotic events and cytokinesis. Plk1 also regulates heat-shock transcription factor 1 (Kim et al. 2005), p53 (Ando et al. 2004) microtubule dynamics (Joukov and De Nicolo 2018), and mitochondrial Ca^2+^ homeostasis (Lee et al. 2016). The tight regulation of Plk1 during the cell cycle is essential to mitotic progression and increased levels of Plk1 have been affiliated with cancer. Increasing evidence indicates that Plk1 overexpression correlates with poor clinical outcomes (Martini et al. 2019), yet the detailed mechanisms of Plk1 regulation have remained unknown. Our data suggests that elevated PLK-1 in early *C. elegans* embryos leads to premature nuclear disassembly and could also account for the observed spindle defects. This finding provides a key link between mitochondria and mitosis by affecting master regulators of mitosis.

## Results and Discussion

### The mitochondrial protein SPD-3 is required for mitotic events

To further determine the functions of the mitochondrial protein SPD-3 during mitosis we characterized the *spd-3(oj35)* mutant, which was previously shown to impact spindle alignment (Dinkelmann et al. 2007). The *spd-3(oj35)* mutant was isolated in an ethyl methanesulfonate (EMS) mutagenesis screen in search of cell division mutants and carries a single cytosine-to-thymidine transition resulting in a leucine-to-phenylalanine change at amino acid 130 (O’Connell et al. 1998; Dinkelmann et al. 2007). The *spd-3(oj35)* strain is a temperature-sensitive, maternal-effect mutant, which is defective in meiosis (Labrador et al. 2013) and mitosis (O’Connell et al. 1998, Dinkelmann et al. 2007). In addition to the *spd-3(oj35)* mutant, 3 additional mutant strains are available (Fig. S1). Two deletion alleles, spd*-3(ok1817)* and *spd-3(tm2969)*, which are null *spd-3* alleles (Labrador et al. 2013). The *spd-3(me85)* mutant strain carries an early stop mutation and has severe meiotic defects impeding the analysis of mitosis (Labrador et al. 2013). Thus, we decided to focus on the effects of *spd-3* mutation in the *spd-3(oj35)* strain.

To characterize the effects on mitotic spindle alignment in *spd-3(oj35)*, we filmed embryos co-expressing mCherry::γ-tubulin and mCherry::histone H2B. Following fertilization in control (WT) embryos, the pronuclear-centrosomal complex moves to the center and rotates to align the mitotic spindle along the anterior/posterior axis (A/P axis) (Fig. S2A). This rotation is impaired in *spd-3(oj35)* embryos, resulting in a mitotic spindle that is misaligned to the A/P axis. Consistent with the previous report (Dinkelmann et al. 2007), we observed 54.5% of spindle misalignment at the metaphase stage in one-cell-stage *spd-3(oj35)* embryos (Fig. S2B). Moreover, the metaphase spindles in the *spd-3(oj35)* embryos are shorter in comparison to control spindles (Fig. 1A, control 14.8 ± 0.6µm (Stdev), *spd-3(oj35)* 12.7 ± 1.4µm). Quantification of the duration of mitosis in one-cell-stage embryos revealed that mitosis is prolonged in the *spd-3(oj35)* mutant embryos. While the *spd-3(oj35)* mutant embryos display a similar duration of the prophase stage of mitosis, which is defined as the time from pronuclear meeting to NEBD (Fig. S2C), the pro-metaphase stage, which is defined as the time from NEBD to anaphase onset, is extended (Fig. 1B). These results suggest that *spd-3(oj35)* does not only affect spindle positioning but also spindle length and the timing of mitotic events.

**Figure 1.**
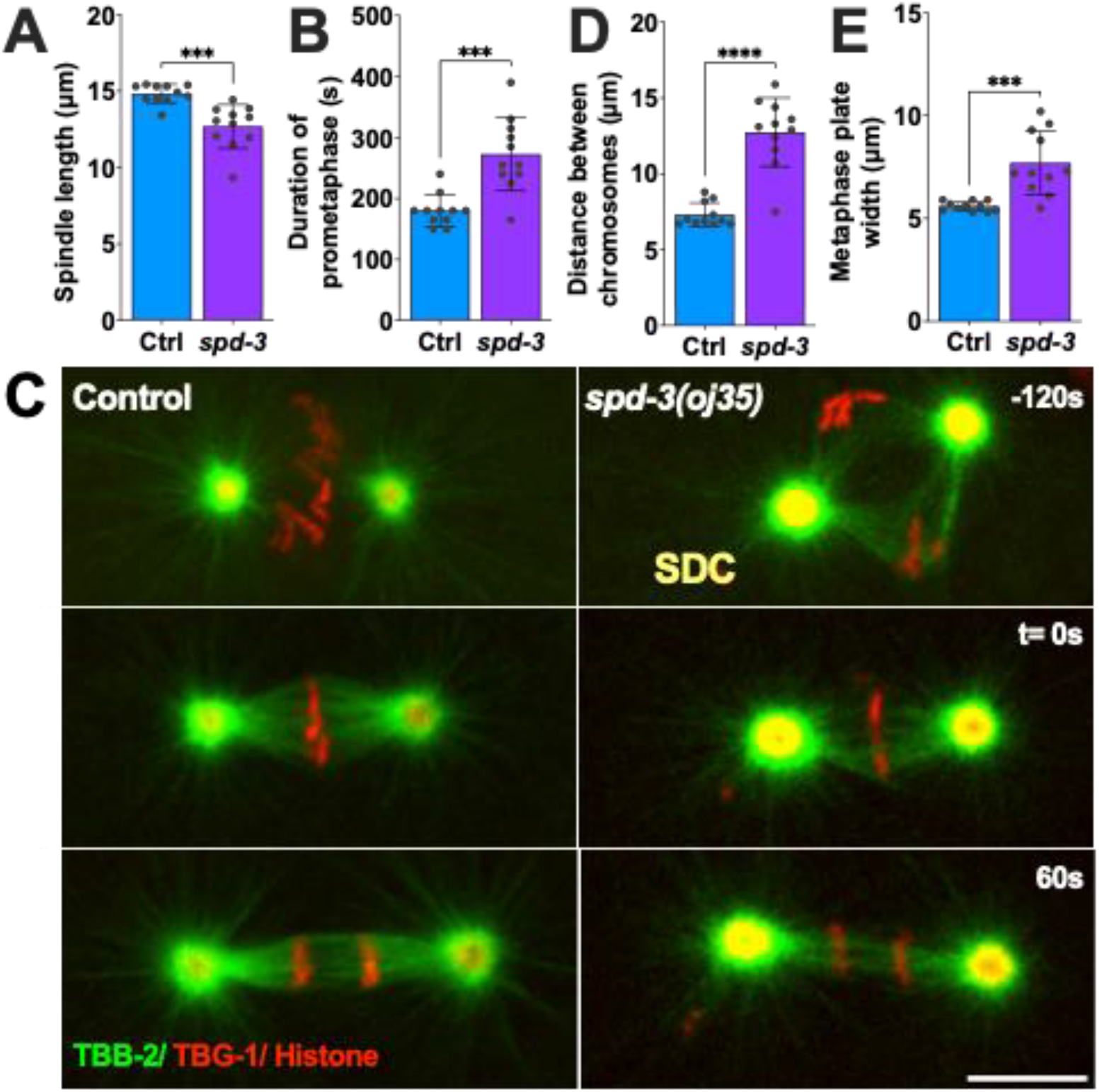
Duration of mitosis, spindle length and chromosome positioning are affected in *spd-3(oj35)* one-cell stage *C. elegans* embryos. *(A)* Graph plotting mitotic spindle length (pole-to-pole distance) at anaphase onset in control (Ctrl, blue bar) and *spd-3(oj35) (spd-3*, purple bar*). (B)* Plot showing prometaphase duration as defined by the time between NEBD and anaphase onset. *(C)* Representative fluorescence confocal images of control (left) and *spd-3(oj35)* (right) embryos expressing GFP::β-tubulin, mCherry::γ-tubulin, and mCherry::histone H2B. SDC indicates the socially distant chromosome stage observed in *spd-3(oj35)* embryos (top right). Scale bar, 10 μm. *(D)* Plot of the distance between chromosomes at the SDC stage. *(E)* Plot of the width of the metaphase plate at anaphase onset. *(B, C)* Times are in seconds relative to anaphase onset (t= 0s). *(A, B, D, E)* Error bars are SD. n = 11 embryos for control and spd-3. The significance of differences between results was determined by two-tailed Student’s t-tests, ***P<0.001, ****P<0.0001.

### The SPD-3 is critical for mitotic chromosome positioning during congression

During our analysis of mitosis in the *spd-3(oj35)* embryos we noticed that the mutant embryos display an additional unusual phenotype during chromosome positioning and alignment. Instead of forming a normal metaphase plate in the center of the embryo, the individual sets of chromosomes (paternal and maternal) are initially positioned on the periphery of the respective nuclei, giving rise to a transient diamond shaped spindle configuration (Fig. 1C, Fig. S2D), before converging to form a metaphase plate. Inspired by the current pandemic we called this abnormal positioning stage the “socially distanced chromosome (SDC) positioning” (Fig. 1C) as the two individual sets of chromosomes, paternal and maternal, showed a large physical distance (Fig. 1D). While the chromosomes eventually form a metaphase plate in the *spd-3(oj35)* mutant, the resulting metaphase plates are significantly wider (Fig. 1E, control: 5.6±0.2 μm, *spd-3(oj35)*: 7.3±1.4 μm) in comparison to control metaphase plates. This observation suggests an additional role for SPD-3 during chromosome congression.

### The mitotic phenotypes of the *spd-3(oj35)* mutant are not caused by changes in energy metabolism

SPD-3 was identified as a mitochondria-localizing protein based on colocalization of SPD-3::GFP and mitochondria stained with MitoTracker Red CMXRos (MTR) (Fig. S1B) and immuno-EM (Dinkelmann et al. 2007, Labrador et al. 2013). We wanted to determine the mechanisms of how the mitochondrial protein SPD-3 affects chromosome positioning. As it was previously suggested that defects in the mitochondrial function in the *spd-3(oj35)* could lead to reduced function of cytoskeletal motors, possibly caused by variations in ATP levels, we measured the metabolic rates in *spd-3(oj35)* embryos by fluorescence lifetime imaging microscopy (FLIM) of NADH autofluorescence (Schneckenburger 1992; Sud et al. 2006). FLIM NADH offers a marker-free readout of the mitochondrial function of cells in their natural microenvironment and allows different pools of NADH to be distinguished within a cell (Blacker et al. 2014). NADH autofluorescence can be used as a marker of cellular redox state and indirectly also of cellular energy metabolism (Schneckenburger 1992; Sud et al. 2006). These measurements show that metabolic rates in *spd-3(oj35)* embryos are increased in comparison to control (Fig. S3). This agrees with previously reported increased ATP levels (Dinkelman et al 2007).

In order to test if elevated (or reduced) ATP could lead to similar mitotic phenotypes we examined the effect of depleting proteins that are part of the electron transfer chain (ETC) in *C. elegans* (Dancy 2015) (Table S1). CLK-1 is required for the biosynthesis of ubiquinone, a carrier in the ETC, and depletion of CLK-1 leads to elevated ATP levels (Braeckman et al. 1999; Dancy 2015). We screened embryos of the null allele *clk-1(qm30)* for defects in spindle and chromosome positioning but could not reproduce any of the observed phenotypes of the *spd-3 (oj35)* mutant in respect to spindle and chromosome positioning. We further analyzed the effects of *isp-1(qm150), ucr-1*(RNAi), *mev-1* (RNAi), or *cco-1* (RNAi), which alter the ETC function resulting in decreased ATP levels (Dillin et al. 2002; Dancy 2015). While we did observe spindle misalignment phenotypes in all examined depletions, we never detected the SDC positioning phenotype in those mitochondrial mutants (Table S1).

In summary, the *spd-3(oj35)* phenotype does not mimic the disruption of mitochondrial genes, suggesting that metabolic perturbations in *spd-3(oj35)* worms may not directly lead to the observed mitotic defects. In agreement with this, also treatment of control embryos with mitochondrial inhibitors (Dinkelman et al 2008) did not reproduce the *spd-3(oj35)* phenotype suggesting that the observed phenotypes are not based on misregulation of ATP levels alone.

### SPD-3 is important for ER morphology

As we observed an effect on the timing from NEBD to anaphase we decided to further investigate the arrangement of ER and nucleus. The ER surrounds and encloses the pronucleus, forming a characteristic bi-membraned nuclear envelope that protects the chromatin. In order to determine the arrangement of the ER in the *spd-3(oj35)* mutant we generated a mutant strain that co-expressed GFP labeled SP12, an ER lumen marker, and mCherry labeled histone. Analysis of this strain revealed that the morphology of the ER is severely affected in the *spd-3(oj35)* mutant (Fig. 2A-C). The ER shows a significant increase in the formation of ER clusters and drastic changes in the morphology surrounding the nucleus (Fig. 2B, C). In control embryos clusters begin to form 238.1±21.9 s before anaphase onset (Fig. 2D) and become enriched around the centrosome and mitotic spindle (Fig. 2B, C). In general, the distribution of ER clusters is asymmetric (Poteryaev et al. 2005), with more clusters localizing in the anterior half of the embryo in the vicinity of the cortex (Fig. 2A). In the *spd-3(oj35)* mutant, excessive ER clusters begin to form earlier, at 367.7±17.5 s before anaphase onset (Fig. 2D), and are found in the cytoplasm, around the nucleus and on the cortex in both the anterior and posterior sides of the *spd-3(oj35)* embryo (Fig. 2A-C). In addition, gaps appear in between centrosome and pronuclei (white arrow) and in between pronuclei (yellow arrow) before anaphase onset in *spd-3(oj35)* embryos. These results show that ER morphology is affected in the early embryonic stage of the s*pd-3(oj35)*, suggesting a potential role for SPD-3 in these processes.

**Figure 2.**
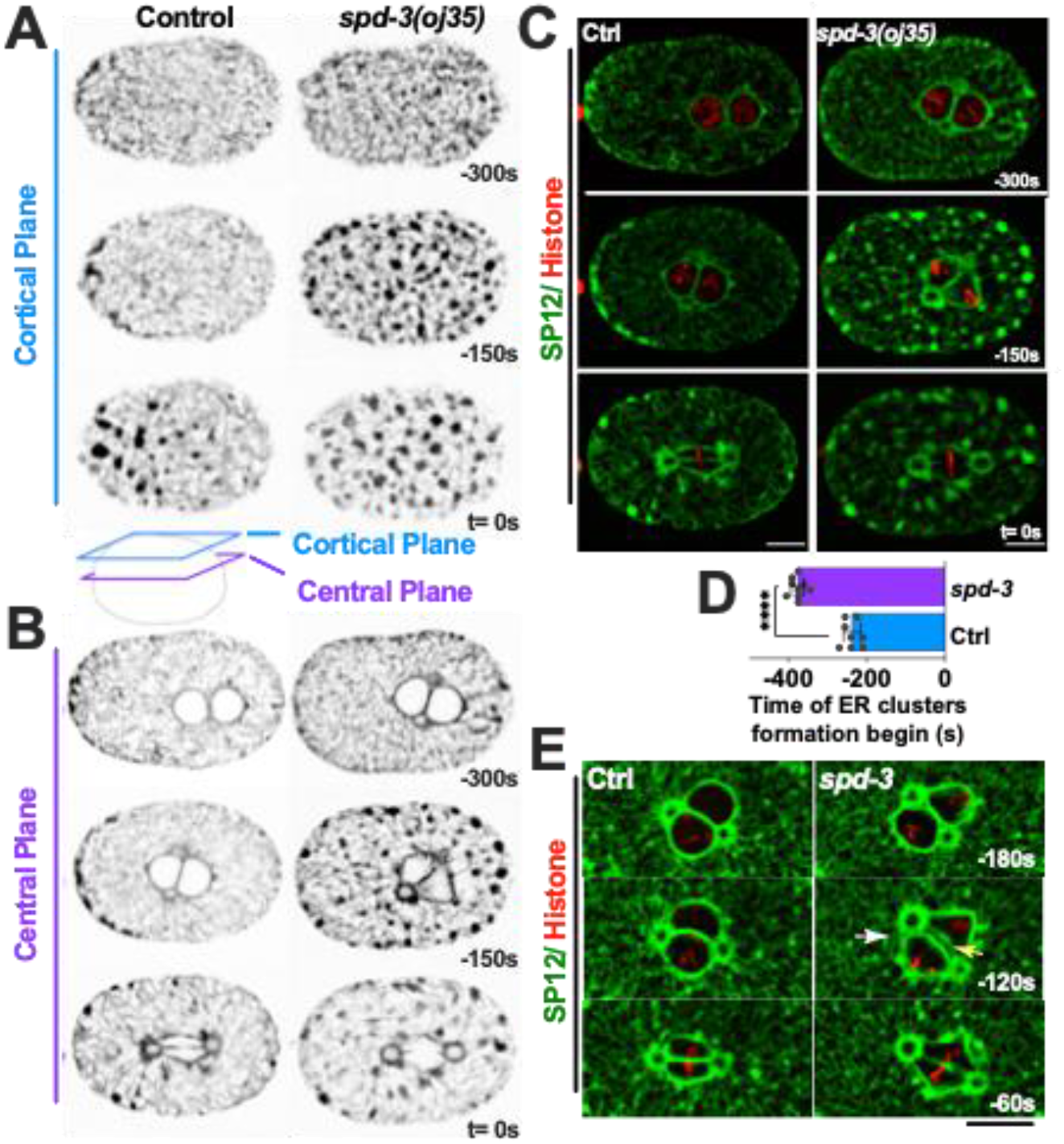
The point mutation of SPD-3 results in abnormal ER morphology. (*A, B*) Spinning-disk confocal images of control (*left*) and *spd-3(oj35) (right)* embryos expressing GFP::SP12 and mCherry::histone H2B. Cartoon indicates the position of a cortical plane just beneath the embryo surface (blue, *A*) and a central plane (purple, *B*) from control and *spd-3(oj35)* are shown. (*C*) Merge of the images shown in A. Control left, *spd-3(oj35)* right (*D*) Graph plotting the time mitotic ER clusters begin to form in the cortical section of embryos expressing GFP::SP12 in mitosis (n = 8 embryos for control and spd-3). Error bars are SD. Significance was determined by t-tests. ****P<0.0001. (*E*) Spinning-disk confocal images of the ER shape surrounding the pronucleus in control (left) and *spd-3(oj35)* (right) embryos coexpressing mCherry::histone H2B and GFP::SP12.. The white arrow indicates a gap appearing between the centrosome and pronuclei (t= −120s). The yellow arrow indicates a gap appearing between the pronuclei (t= −120s). (*A-E*) Times are in seconds relative to anaphase onset (t= 0s). Scale bars, 10 μm.

In order to determine the potential effects of ER morphology on spindle and chromosome positioning, we analyzed a number of ER mutants that show drastic changes in ER morphology, in particular the ratio of clusters to tubes. The RAB-5 and YOP-1/RET-1 proteins are important for regulating the ER morphology during mitosis and controlling the kinetics of nuclear envelope disassembly (Audhya et al. 2007). The endosomal Rab-type GTPase, RAB-5, plays a role in the homotypic fusion of ER membranes (Audhya et al. 2007). The YOP-1, homologs of DP1/NogoA, localize to the ER and play a functionally redundant role in generating tubular morphology in the ER (Audhya et al. 2007). In YOP-1 and RAB-5 depleted embryos, there are fewer thick ER tubules that are poorly organized and no mitotic ER clusters form. We treated *spd-3(oj35)* worms with *rab-5* (RNAi) (data not shown) and *yop-1* (RNAi) with the aim to reduce the number of ER clusters in the cytoplasm. While we did observe fewer and smaller ER clusters in the *spd-3(oj35)* mutant after *rab-5* and *yop-1* depletion, the SDC positioning phenotype was still present (Fig. S4).

Next, as ER dynamics and morphology changes can also be induced by ER stress, we tested whether the ER stress-unfolded protein response (UPR^ER^) is changed in the *spd-3(oj35)* mutant. During *C. elegans* UPR^ER^ two homologs of the ER-resident heat-shock protein BiP, HSP-3 and HSP-4, are essential for the formation of sheet-like ER structures (Poteryaev et al. 2005). Depletion of HSP-3 leads to upregulation of the *hsp-4* gene in *C. elegans* (Kapulkin et al. 2005). To examine the UPR^ER^ in *spd-3(oj35)*, we imaged worms expressing the *zcIs4* transgene, a transcriptional fusion of the *hsp-4* promoter to GFP, that can be used to quantify HSP-4 expression levels (Calfon et al. 2002). The zcIs*4* reporter is strongly upregulated in response to treatments inducing ER stress and can thus function as a read out for UPR^ER^ activity (Calfon et al. 2002). We found an overall lower HSP-4 expression level in *spd-3(oj35)* mutant worms by quantification of fluorescence levels, indicating an approximately 11 fold decrease in *hsp-4::gfp* expression (Fig. S5A, B). This result suggested that changes in ER morphology could be induced by blocking UPR^ER^ leading to low HSP-4 expression levels in the *spd-3(oj35)* mutant. To further determine whether the mitotic defects could be rescued by increasing HSP-4, we treated *spd-3(oj35)* mutant with *hsp-3* (RNAi) to increase the endogenous *hsp-4* transcription (Kapulkin et al. 2005). However, the chromosome positioning and spindle defects in the *spd-3(oj35)* mutant are not prevented by *hsp-3* (RNAi) (data not shown). In control embryos *hsp-3* (RNAi) does not affect ER dynamics in the early embryo. In contrast, when HSP-4 function is reduced by RNAi, the ER accumulates in multiple foci, and the ER morphology is distinctly disrupted. However, we did not observe any chromosome positioning and spindle defects in HSP-4 depleted embryos (data not shown). Taken together, our results suggest that perturbation of the mitochondrial protein SPD-3 affects the morphology of the ER and that this might be induced by the UPR^ER^ regulatory system. However, the structural changes of the ER are most likely not responsible for the observed chromosome positioning and spindle alignment defects in the *spd-3(oj35)* mutant embryos.

### SPD-3 is critical for nuclear envelope structure and dynamics

Our analysis of the ER morphology revealed a significant change in the shape of the pronucleus in *spd-3(oj35)* mutant. As the ER and the nucleus are intimately connected, we generated strains that carry either NPP-22::mNeonGreen (NPP-22::mNG) (Mauro et al. 2022), a conserved transmembrane nucleoporin, or GFP tagged lamin (Link et al. 2018), which enabled us to analyze the shape and organization of the nuclear envelope during mitosis. Analysis of these strains by light-microscopy showed that the nuclear shape is significantly altered during prometaphase in *spd-3(oj35)*, consistent with the observed changes in ER morphology surrounding the spindle (Fig. 3A). Analysis of the lamin GFP in the *spd-3(oj35)* mutant background showed that in comparison to control embryos lamin is disassembled prematurely, approximately 96.5s earlier in comparison to control embryos (Fig. 3B, C). In addition, the lamin of the paternal pronucleus (164.8±52.8s) disappears earlier than the lamin on the maternal pronucleus (120.8±38.4s) (Fig. 3D). This indicates that the structural integrity of the nucleus is impaired in the *spd-3(oj35)* mutant and that this could potentially affect the positioning of the spindle as well as chromosomes, possibly by destabilizing nuclear membrane attachments to the centrosome, microtubules and chromosomes.

**Figure 3.**
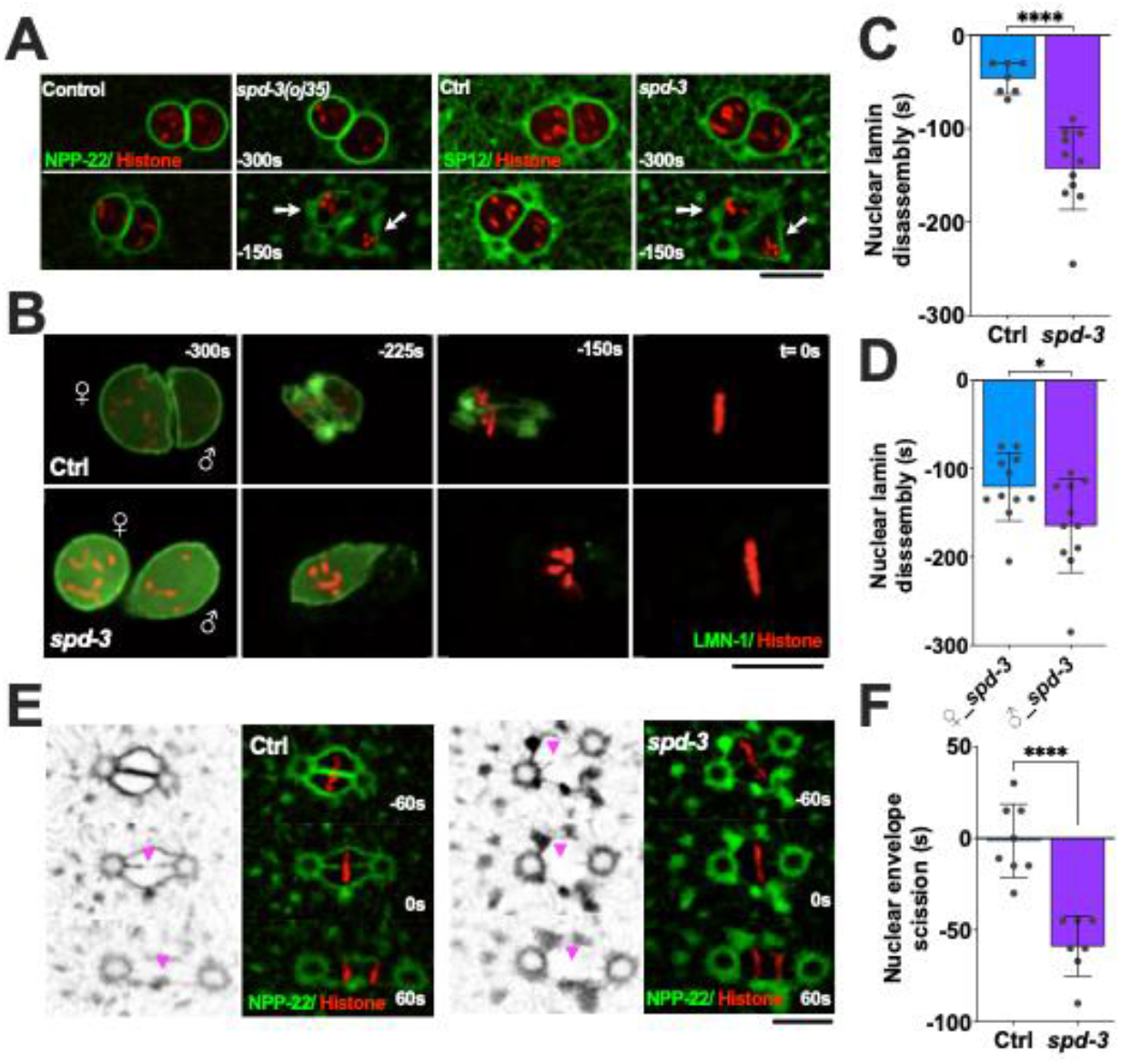
SPD-3 is involved in structural integrity and disassembly of the nuclear envelope during mitosis. (*A*) Spinning-disk confocal images of nuclear morphology in control (Ctrl) and *spd-3(oj35)* (*spd-3*) embryos coexpressing mCherry::histone H2B and NPP-22::mNG (*left*) or GFP::SP12 (*right*). The white arrows indicate the abnormal nuclear shape in *spd-3(oj35)*. (*B*) Time-lapse sequences of control (*left*) and *spd-3(oj35)* (*right*) embryos expressing GFP::LMN-1 and mCherry::histone H2B. ♀ indicates female pronucleus and ♂ indicates male pronucleus. Lamin disassembles prematurely and asymmetrically in *spd-3(oj35)* embryos. (*C*) Plot of the timing of complete nuclear lamin disappearance in embryos co-expressing GFP::LMN-1 and mCherry::histone H2B in control (n=7) and *spd-3(oj35)* (n=11). (*D*) Plot of the timing of complete lamin disassembly in the female (*left*, n=11) and male pronucleus (*right*, n=11) in *spd-3(oj35)* embryos. (*E*) Fluorescence confocal images of control and *spd-3(oj35)* embryos expressing NPP-22::mNG and mCherry::histone H2B. Magenta arrowheads mark the site of nuclear envelope scission in control (*left*) and *spd-3(oj35)* (*right*). (*F*) Graph plotting time of nuclear envelope scission in control (n=8) and *spd-3(oj35)* (n=7) embryos. (*A-F*) Times are in seconds relative to anaphase onset (t= 0s). Scale bars, 10 μm. (*C, D, F*) Error bars are SD. The significance of the difference between strains was determined by t-tests. *P<0.05, ****P<0.0001.

### Early nuclear envelope scission in prometaphase

The timely disassembly of the nuclear lamina is required for chromosome alignment and NE scission to allow parental chromosomes to merge on the metaphase plate in *C. elegans* (Rahman et al. 2015; Chase et al. 2000; Velez-Aguilera et al. 2020). We next examined whether NE scission changes between the juxtaposed pronuclei by labeling of NE (NPP-22::mNG) or ER (GFP::SP12). The NE scission event occurs less than 3 s (1.4-2.5 s) before anaphase onset in control embryos and the forming membrane gap is visible (Fig. 3E, F, Fig. S6A-C). In contrast, the NE scission event occurs early, 40-60 s (37.5-58.9 s) before anaphase onset, in *spd-3(oj35)* embryos.

In control embryos the reformation of new NE around the segregated chromosomes begins approximately 86.7±20.9 s after anaphase onset and the new NE is completed after 161.7±27.8 s (Fig. S6D, E). The reassembly of the NE is slightly delayed in *spd-3(oj35)* embryos as it is initiated 101.7±18.0 s and completed 180.0±29.0 S after anaphase onset (Fig. S6D, E). Combined these results suggest that SPD-3 is critical for the dynamics of NE disassembly during the first mitotic division in *C. elegans*.

### Spindle and chromosome positioning defects in *spd-3(oj35)* are caused by PLK-1 overexpression

Nuclear disassembly is triggered by phosphorylation of lamin at multiple residues in the head and tail domain by PLK-1 in *C. elegans* (Chase et al. 2000; Rahman et al. 2015; Velez-Aguilera et al. 2020). Therefore, we generated a strain expressing PLK-1::GFP in *spd-3(oj35)* mutant to monitor the localization and expression of PLK-1. To our surprise, this strain revealed excessive amounts of PLK-1 in the *spd-3(oj35)* mutant localizing to centrosomes, the nuclear envelope, nucleoplasm, and chromosomes (Fig. 4A, C). We further confirmed that PLK-1 is increased by around 6-fold in the *spd-3(oj35)* mutant compared to N2 animals by immunoblot (Fig. 4B).

**Figure 4.**
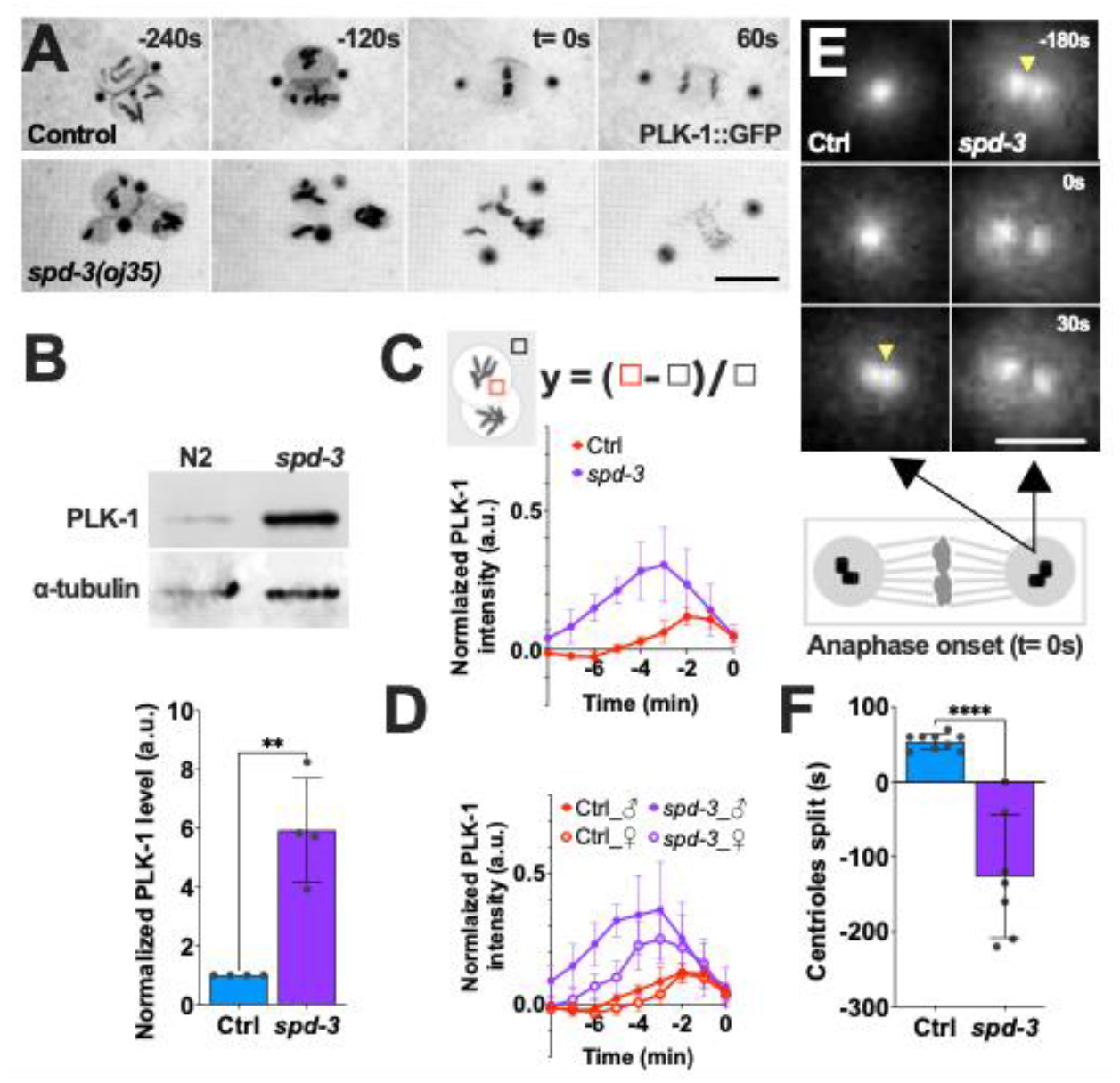
PLK-1 overexpression in the *spd-3(oj35)* mutant. (*A*) Time-lapse sequences of embryos expressing PLK-1::GFP in control (Ctrl, *top*) and *spd-3(oj35)(spd-3, bottom)*. Scale bar, 10 μm. (*B*) Top: Western of PLK-1 protein in N2 (n=4) and *spd-3(oj35)* (n=4) worms, bottom: Quantification of PLK-1 expression level. (*C, D*) Quantification of normalized PLK-1::GFP intensity in the nucleoplasm of embryos measured from time-lapse sequences as in (*A*). (*C*) The value of PLK-1::GFP intensity in control (red, n=6) and *spd-3(oj35)* (purple, n=6). (*D*) The value of PLK-1::GFP intensity in the nucleoplasm of male and female pronuclei. Solid circles indicate the male pronucleus (n=6), empty circles indicate the female pronucleus (n=6). Cartoon showing the method used for quantification of the PLK-1::GFP fluorescence in nucleoplasm. Briefly a fixed-size box is drawn in nucleoplasm (red box) and in cytoplasm (black box) as background at each time point. The normalized PLK-1 level is calculated as [(integrated intensity in red box – integrated intensity in black box)/ integrated intensity in black box]. ♀ indicates female pronucleus and ♂ indicates male pronucleus. (*E*) Spinning-disk confocal images of the centrosome in control (*left*) and *spd-3(oj35)* (*right*) embryos expressing PLK-1::GFP. Yellow arrowheads indicate the time of centrioles splitting. Schematic showing that each centrosome contains a pair of centrioles. Black arrows indicate a centrosome. Scale bar, 2 μm. (*F*) Graph plotting the time of detectable centriole pair splitting in control (*left*, n=6 centrosomes) and *spd-3(oj35)* (*right*, n=6 centrosomes). (*A, C, D, E, F*) Times are in seconds relative to anaphase onset (t= 0). (*B, C, D, F*) Error bars are SD. The significance of the difference between strains was determined by t-tests. **P<0.01, ****P<0.0001.

In *C. elegans*, PLK-1 is recruited to the nuclear pore complexes (NPC) by nucleoporins NPP-1, NPP-4, and NPP-11 prior to NEBD where the importins α/β IMA-2 and IMB-1 promote the nuclear import of PLK-1, where it phosphorylates nuclear lamin (Martino et al. 2017). To determine the time point of PLK-1 import into the nucleus, we quantified the nuclear fluorescence of PLK-1::GFP in nucleoplasm during mitosis. The intensity of PLK-1::GFP reaches a peak about 2 minutes prior to anaphase onset in both pronuclei of control embryos (Figure 4C). This peak is shifted in *spd-3(oj35)* embryos to about 3 minutes prior to anaphase onset. In addition we detected an overall increased fluorescence of PLK-1::GFP in the paternal pronucleus relative to the maternal pronucleus (Fig. 4D). This result is consistent with the observed premature, faster lamin disassembly in the paternal pronucleus (Fig. 3D). In addition to the nucleoplasm, more PLK-1::GFP localizes to centrosomes. Centrosomes consist of a centriole pair surrounded by pericentriolar material (PCM) that nucleates and anchors microtubules (MTs) in *C. elegans*. In general, the centriole pair splits 53.8±10.5 s after anaphase onset during mitosis (Fig. 4E, F). Surprisingly, the centriole pair splits 126.4±82.1 s before anaphase onset in *spd-3(oj35)* embryos (Fig. 4E, F). This result is consistent with overexpression and inhibition of Plk1 affecting the centrioles disengagement (Lončarek et al. 2010; Schöckel et al. 2011; Fu et al. 2015). This data indicates that overexpression of PLK-1 leads to premature centrioles splitting independent of the cell cycle regulation.

To determine whether reducing PLK-1 in the nucleoplasm can avoid the socially distanced chromosomes and rescue spindle misalignment, we exposed the *spd-3(oj35)* worms to *ima-2* (RNAi) to prevent nuclear import of PLK-1. In control embryos *ima-2* (RNAi) prevents the nuclear import of PLK-1 and this its localization to kinetochores and chromatin (Martino et al. 2017) (Fig. S7A). In addition, previous work showed that *ima-2* (RNAi) is essential for spindle assembly and nuclear envelope formation in the *C. elegans* embryo (Askjaer et al 2002). In *spd-3(oj35)* mutant embryos, PLK-1::GFP still localizes to nucleoplasm and chromatin after *ima-2 (RNAi)* in embryos (Fig. S7A), however the amount of PLK-1 is reduced and the peak of PLK-1 fluorescence is shifted to about 2 minutes prior to anaphase onset, similar to control embryos (Fig. S7B). Interestingly, the SDC and spindle misalignment are rescued by *ima-2* (RNAi) supporting the hypothesis that both phenotypes are induced by elevated amounts of PLK-1 in the nucleus (Fig. S7C, D). To test this hypothesis, we exposed the *spd-3(oj35)* worms to short treatments of *plk-1* (RNAi) to reduce the elevated levels of PLK-1 to approximately wild-type levels (Fig 5A, B). Reduction of the PLK-1 overexpression does indeed rescue the socially distanced chromosome positioning (Fig. 5C). The spindle positioning defects seemed slightly reduced but this reduction was not significant (Fig. S7E). This supports our hypothesis that PLK-1 overexpression in the *spd-3(oj35)* mutant affects the nuclear integrity by inducing premature and asymmetric nuclear envelope disassembly and that this most likely affects proper chromosome and spindle positioning.

**Figure 5.**
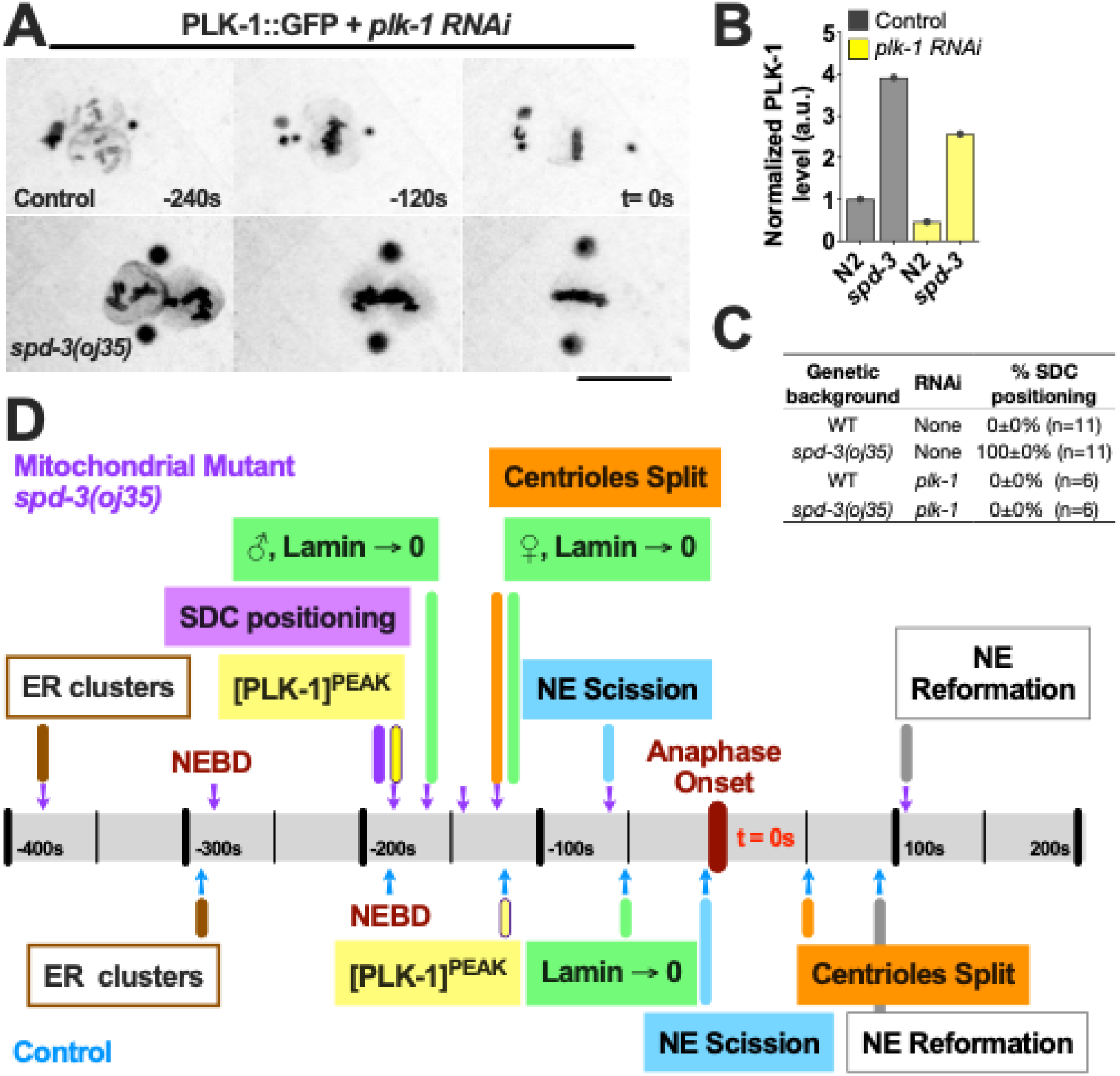
Mitotic phenotypes in the *spd-3(oj35)* embryos result from PLK-1 overexpression. (*A*) Time-lapse sequences of embryos expressing PLK-1::GFP in control (Ctrl, *top*) and *spd-3(oj35)* (*spd-3, bottom*) after *plk-1 (RNAi)* were filmed using a spinning-disk confocal. Scale bar, 10 μm. (*B*) Graph of the quantification of PLK-1 expression level in lysate from N2 and *spd-3(oj35)* without (gray bars, n=1 for each strain) and with *plk-1 (RNAi)* (yellow bars, n=1 for each strain) as determined by western blot. (*C*) Plot of the percentage of socially distanced chromosome (SDC) positioning in control and *spd-3(oj35)* without (gray bars, n=6 for each strain) and with *plk-1 (RNAi)* (yellow bars, n=6 for each strain) treatment. (*D*) Schematic showing the timeline of mitotic events in control and *spd-3(oj35)*. (*A, D*) Times are in seconds relative to anaphase onset (t= 0).

To determine if the abnormal ER morphology is also triggered by PLK-1 overexpression we examined the ER structure after *plk-1* (RNAi). We found that the ER morphology and distribution of ER clusters are not significantly different between control and *plk-1* (RNAi) in *spd-3(oj35)* embryos, suggesting that the changes in ER morphology are not induced by elevated PLK-1 levels (Fig. S8A, B). Our finding is thus consistent with the result that the removal of ER clusters cannot rescue socially distanced chromosomes (Fig. S4). Taken together we show that defects in SPD-3 cause a spindle positioning defect and an unusual chromosome positioning, socially distanced chromosomes (SDC), phenotype during mitosis in *spd-3(oj35)* embryos. The fact that we did not observe comparable phenotypes in embryos defective in ATP production, suggests that mitochondrial ETC pathways are not involved in this process. The abnormal spindle and chromosome positioning are accompanied by altered ER and NE morphology, and premature and asymmetric nuclear lamin disassembly in prometaphase (Fig. 5D).

Our results revealed an increase in PLK-1 levels in the *spd-3(oj35)* embryos. As PLK-1 plays a direct role in lamin disassembly, by phosphorylating lamin and thus targeting it for degradation (Velez-Aguilera et al. 2020), this suggests that the elevated levels of PLK-1 are driving the premature lamin disassembly in the *spd-3(oj35)* mutant. In addition, our data showed that the paternal pronucleus disassembled several seconds before the maternal pronucleus. As PLK-1 localizes to centrosomes, which are initially associated with the male pronucleus this indicates that the asymmetric disassembly of the two pronuclei is also linked to excessive levels of PLK-1. We propose that the chromosome alignment defects and premature lamin disassembly, the previously described spindle positioning defect (Dinkelmann et al, 2007), and the reported reduction in mobility of SUN-1 aggregates and pairing-center binding proteins on the nuclear envelope in *spd-3(me85)* mutant (Labrador et al 2013) could be linked to the elevated PLK-1 levels in the *spd-3(oj35)* mutant.

The linker of nucleoskeleton and cytoskeleton (LINC) complex connects the nuclear lamina to cytoplasmic MTs and also contributes to centrosome attachment to the outer nuclear membrane (Cain et al. 2018). The phosphorylation of SUN1, a component of the LINC complex by Plk1 and CDK1, inhibits SUN1 interaction with the nuclear lamina (Labella et al. 2011; Patel et al. 2014). It is possible that more PLK-1 in the nucleoplasm leads to a destabilization of the outer nuclear membrane attachment to the centrosome by phosphorylation of SUN-1 and premature lamin disassembly and that this could affect spindle positioning. In addition, elevated levels of PLK-1 on the chromosomes could also affect kinetochore assembly and function and thus result in chromosome positioning defects. Lastly, we noticed that the male pronucleus expands before the pronuclear meeting (Fig. S9), the PCM increases in size and microtubule nucleation is increased in *spd-3(oj35)*, which could all be induced by increased PLK-1 levels (Fig. S10).

Besides mitosis, PLK-1 also plays roles in mitochondrial Ca^2+^ homeostasis and ATP production (Lee et al. 2016). It has been shown that mitochondrial Ca^2+^ can positively regulate the activities of the tricarboxylic acid (TCA) cycle and electron transport chain components, stimulating ATP production (Shanmughapriya et al. 2015). The mitochondrial rho GTPase Miro controls mitochondrial Ca^2+^ homeostasis at the ER-mitochondria contact sites, and its activity is regulated by Polo kinase in *Drosophila* (Lee et al. 2016). Since *spd-3(oj35)* has excessive amounts of PLK-1, elevated ATP levels could be triggered by increased Miro activation affecting Ca^2+^ import into mitochondria. It would be interesting to further examine the interaction between ER and mitochondria and mitochondrial Ca^2+^ homeostasis in *C. elegans*.

The remaining question is how the mitochondrial protein SPD-3 affects or regulates PLK-1 levels during mitosis in *C. elegans*. At this point we can only speculate, but possible mechanisms include i) regulation of mRNA levels (Martin and Strebhardt 2006), ii) mRNA translation (Tanenbaum et al. 2015), iii) post-translational modifications (Macůrek et al. 2008), or iv) protein stability (Cordeiro et al. 2020). Further detailed analysis will be necessary to determine the mechanisms and pathways utilized by SPD-3 that led to increased PLK-1 levels.

In summary, our work provides a novel link between mitochondria, nuclear envelope dynamics and chromosome positioning by increasing the amount of a key mitotic regulator, PLK-1. This finding does not only provide a new role of the mitochondrial protein SPD-3 during mitosis in *C. elegans* but may also have further implications in the context of cancers or age-related diseases and infertility as it provides a novel link between mitochondria and mitosis.

## Materials and methods

### C. elegans strains and RNA-mediated interference

All *C. elegans* strains were cultured at 16°C before imaging and experiments (Brenner 1974). *C. elegans* strains and RNAi feeding clones are listed in the Supplemental Material. L4 or young adult hermaphrodites were fed with RNAi bacteria and incubated for 6-48 h at room temperature (23°C) before dissection to obtain embryos for filming (for details, see the Supplemental Material).

### Live cell imaging

For live imaging of embryos, worms were dissected on glass coverslips in M9 buffer and then mounted on 2% agar pads. Imaging was conducted at 23°C (room temperature) and carried out on a on a 3i VIVO spinning-disc confocal microscope (Z1; Carl Zeiss) equipped with 488 nm and 561 nm diode laster and a Hamamatsu ORCA-Flash4.0 scientific CMOS camera (Hamamatsu) for detection. Acquisition parameters were controlled using the Sildebook 6.0 software (3i - Intelligent Imaging). For details, see the Supplemental Material.

## Acknowledgments

We thank the *Caenorhabditis* Genetics Center, which is funded by the National Institutes of Health (NIH) National Center for Research Resources (NCRR), Jessica L. Feldman (Stanford University), Shirin Bahmanyar (Yale University), and Verena Jantsch-Plunger (University of Vienna) for strains; Orna Cohen-Fix (NIH) for RNAi feeding clones; Monica Gotta (University of Geneva) for PLK-1 antibody; Thomas Müller-Reichert (Technische Universität Dresden), Kevin F. O’Connell (NIH), and Shirin Bahmanyar (Yale University) for critical comments and discussions; Horst Wallrabe for the 2-photon FLIM microscopy training (The W.M. Keck Center for Cellular Imaging, University of Virginia). This work was supported by grant from the NIH NIGMS 1R01GM144668.

---

**Table S1.** Strains and RNAi feeding clones for the electron transfer chain (ETC) pathway and sorting and assembly machinery (SAM) complex in mitochondria, related to Figure 1.

| Gene | % SDC positioning | % Spindle misalignment | % Abnormal nuclear shape |
| --- | --- | --- | --- |
| Control | 0 (n=11) | 0 (n=11) | 0 (n=11) |
| *spd-3(oj35)* | 100 (n=11) | 55.6 (n=11) | 100 (n=11) |
| *ucr-1 RNAi* | 0 (n=3) | 33.3 (n=3) | 0 (n=3) |
| *mev-1 RNAi* | 0 (n=2) | 50.0 (n=2) | 0 (n=3) |
| *isp-1(qm150)* | 0 (n=10) | 20.0 (n=10) | N.A. |
| *cco-1 RNAi* | 0 (n=5) | 40.0 (n=5) | 0 (n=3) |
| *clk-1(qm30)* | 0 (n=10) | 0 (n=10) | N.A. |

**Figure S1.**
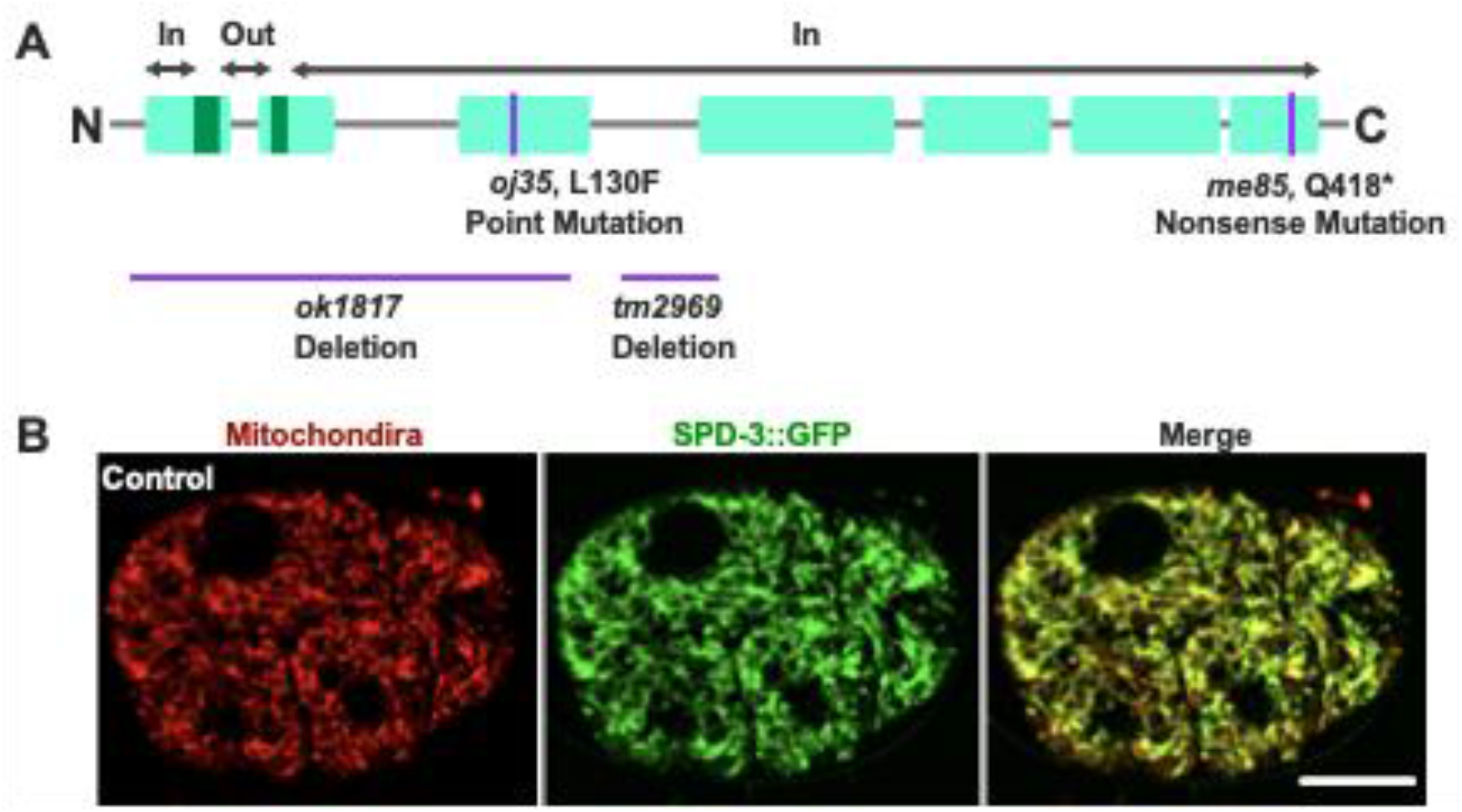
SPD-3 localizes to mitochondria, related to Figure 1. (**A**) Diagram of the *spd-3* gene indicating the position of the *oj35* and *me85* mutation, and two deletion alleles, *ok1817* and *tm2929*. (**B**) SPD-3::GFP colocalizes with mitochondria stained by the mitochondrial dye MitoTracker CMXRosetta. Scale bar, 10 μm.

**Figure S2.**
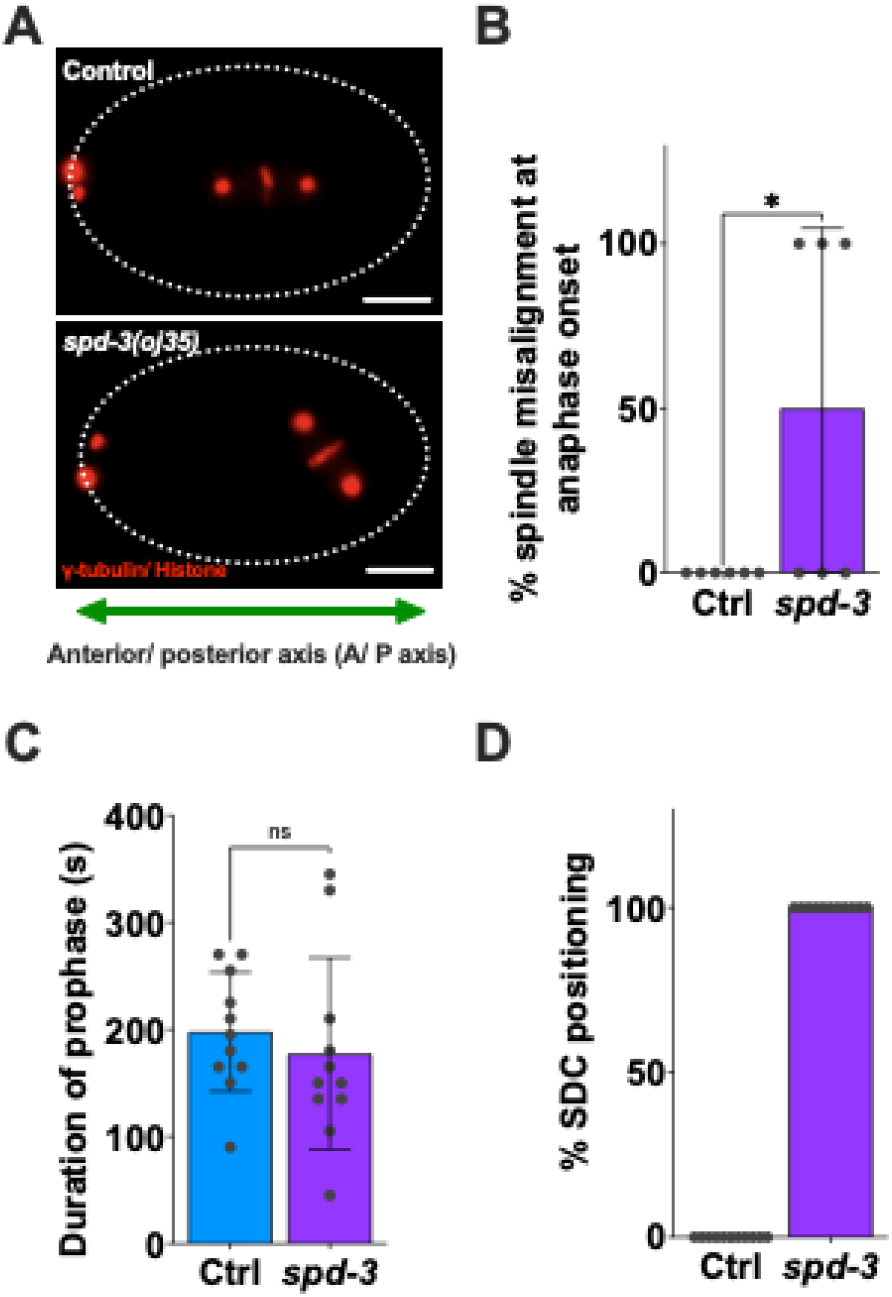
*spd-3(oj35)* is defective in mitotic spindle alignment and chromosome positioning, related to Figure 1. (**A**) Images of embryos coexpressing mCherry::histone H2B and mCherry::γ-tubulin in control (**top**, Ctrl) and *spd-3(oj35)* (**bottom**, *spd-3*). In control embryos, the mitotic spindle is aligned along the anterior/posterior axis (A/P axis). The mitotic spindle failed to misalign to the A/P axis in *spd-3(j35)* embryos. Scale bars, 10 μm. (**B**) Plot of the percentage of spindle misalignment in control (n=11) and *spd-3(oj35)* (n=11) embryos at anaphase onset. (**C**) Plot of the duration of mitotic prophase. Duration of prophase is defined by the time between pronuclei meeting and NEBD (n=11 embryos for control and *spd-3*). Error bars are SD. The significance of the difference between strains was determined by t-tests. ns indicates no significant difference. (**D**) Graph plotting the percentage of socially distanced chromosomes during mitosis (n>11 embryos for control and *spd-3*).

**Figure S3.**
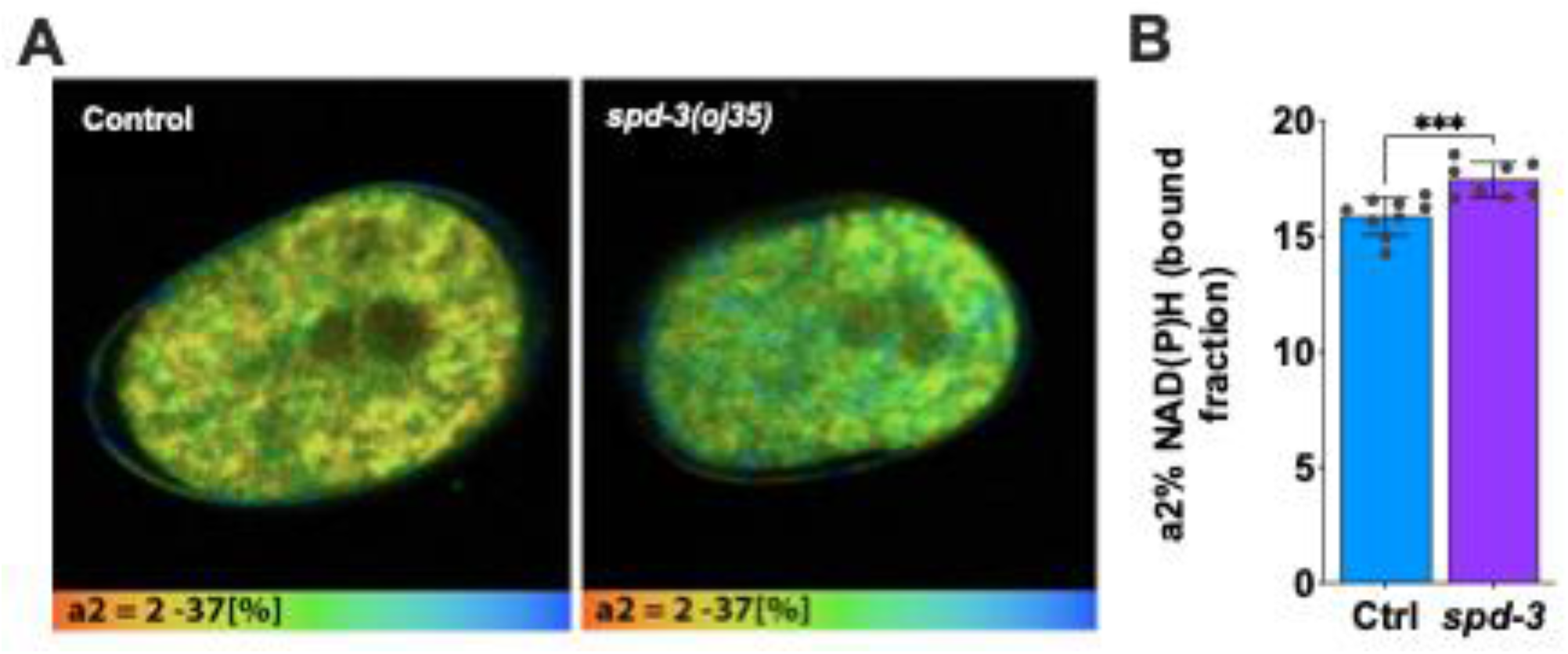
FLIM NAD(P)H imaging shows increased metabolic rates in *spd-3(oj35)* embryos, related to Figure 1. (**A**) Images showing color-coded values of NAD(P)H bound fraction in control (Ctrl) and *spd-3(oj35)* (*spd-3*) one cell stage *C. elegans* embryo. Bottom gradient indicates the level of bound NAD(P)H reaching from 2-37%. (**B**) Bar chart graph comparing mean values for NAD(P)H bound fraction (a2%) in tested embryos. Mean pixel value was calculated from 2-3 image acquisitions per embryo. n=9 for control and n=8 for *spd-3(oj35)*.

**Figure S4.**
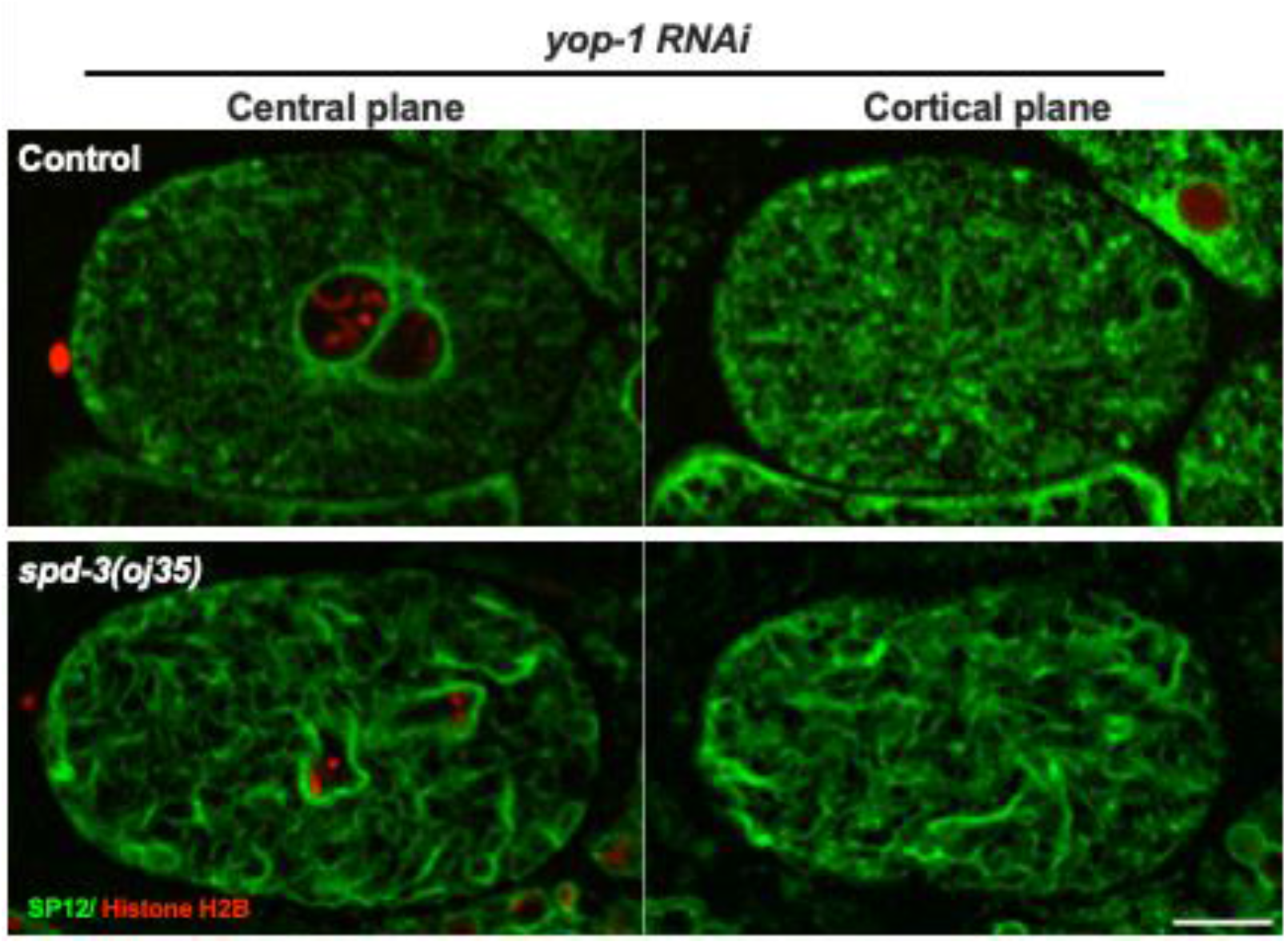
Socially distanced chromosome positioning is not rescued by the removal of ER clusters, related to Figure 2. Spinning-disk confocal images of embryos in control (**top**) and *spd-3(oj35)* (**bottom**) with *yop-1 (RNAi)*. Representative images of a central (**left**) and cortical plane (**right**) are shown. Embryos express the GFP-tagged ER marker SP12 and histone H2B labeled with mCherry. Scale bar, 10 μm.

**Figure S5.**
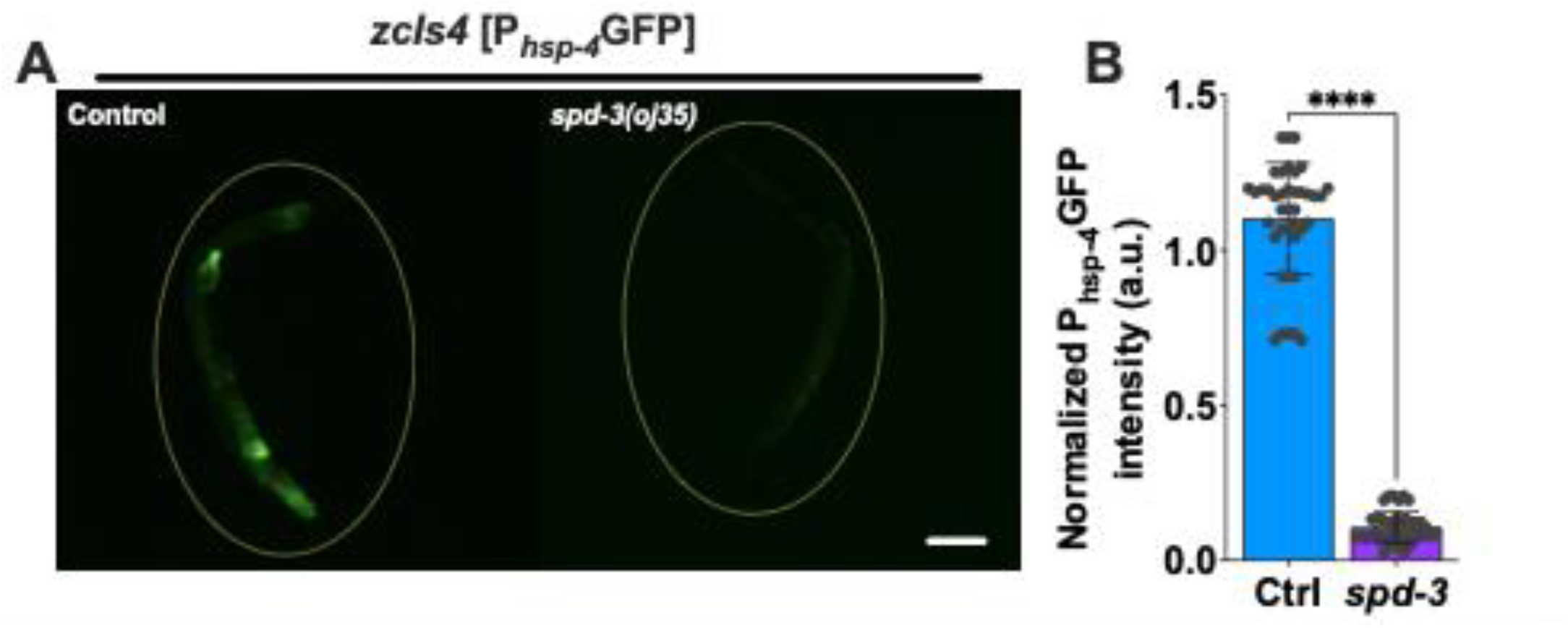
Reduction of *hsp-4::gfp* in *spd-3(oj35)* adult animals, related to Figure 2. (**A**) Fluorescence images of control (Ctrl) and *spd-3(oj35)* (*spd-3*) adult animals carrying the P_*hsp-4*_∷GFP (*zcIs4*) transgene. Scale bar, 100 μm. (**B**) Graph plotting the normalized overall fluorescence of *zcIs4* in control (n=40) and *spd-3(oj35)* (n=40). Error bars are SD. The significance of the difference between strains was determined by t-tests. ****P<0.0001.

**Figure S6.**
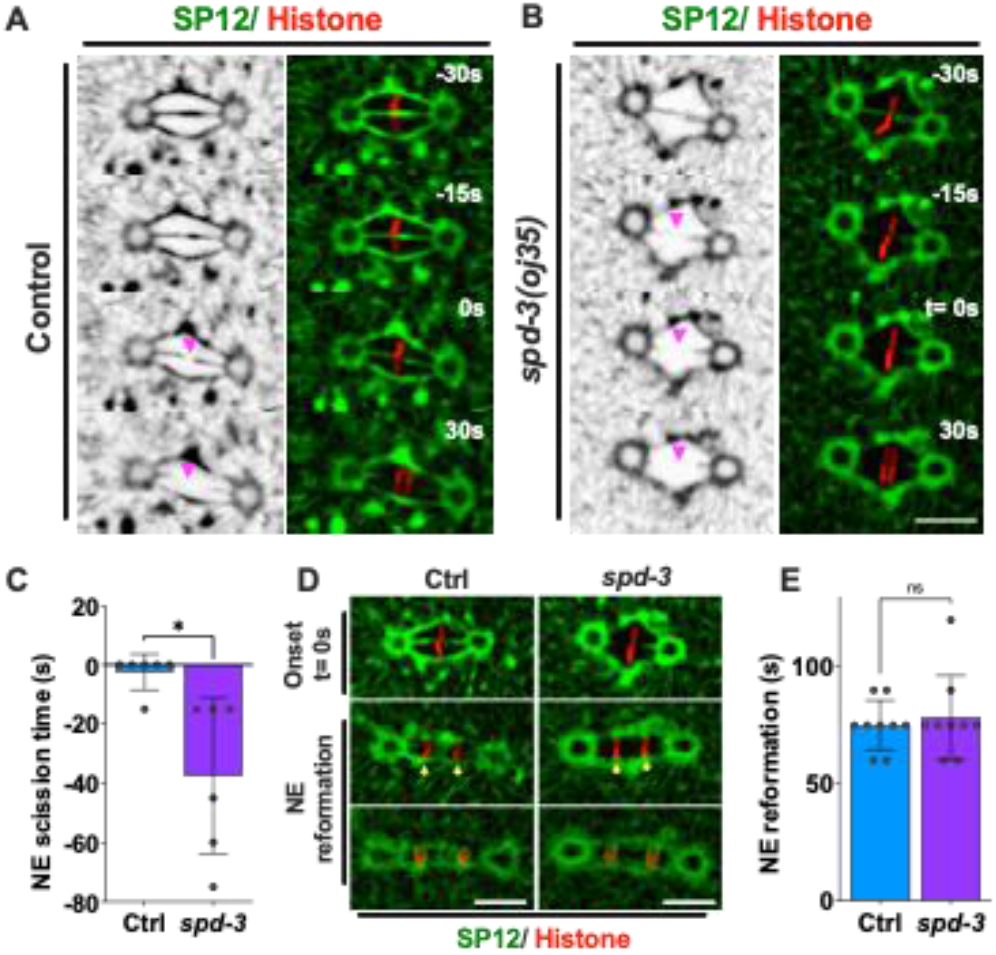
SPD-3 is required for nuclear envelope disassembly and assembly, related to Figure 2. (**A, B**) Fluorescence confocal images of control (Ctrl) and *spd-3(oj35) (spd-3)* embryos expressing GFP::SP12 and mCherry::histone H2B. The left panel shows SP12 alone, the right panel shows a merge of SP12 and Histone. Magenta arrowheads mark the site of nuclear envelope scission in control (**A**) and *spd-3(oj35)* (**B**). (**C**) Graph plotting time of nuclear envelope scission in control (**left**, n=7) and *spd-3(oj35)* (**right**, n=7). (**D**) Fluorescence confocal images of control and *spd-3(oj35)* embryos expressing GFP::SP12 and mCherry::histone H2B. Yellow arrowheads mark the site of nuclear envelope reformation in control (**top**) and *spd-3(oj35)* (**bottom**). (**E**) Graph plotting time of nuclear envelope reformation in control (left, n=9) and *spd-3(oj35)* (right, n=9). (**A, B, D**) Scale bars, 10 μm. (**A-E**) Times are in seconds relative to anaphase onset (t= 0). (**C, E**) Error bars are SD. The significance of the difference between strains was determined by t-tests. ns indicates no significant difference. *P<0.05.

**Figure S7.**
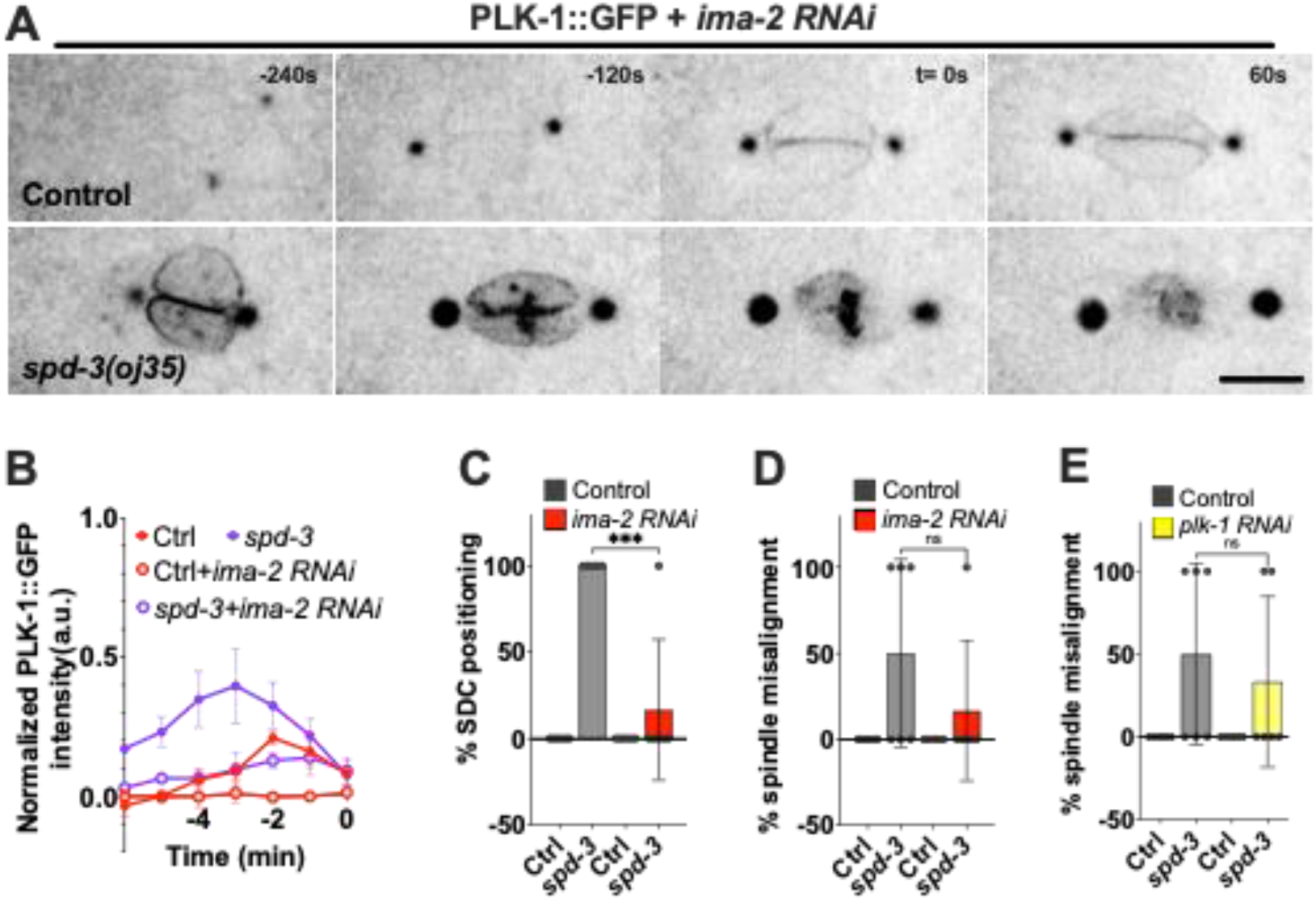
Reduction of PLK-1 in nucleoplasm can prevent the socially distanced chromosome positioning and spindle misalignment, related to Figure 4. (**A**) Time-lapse sequences of embryos expressing GFP::PLK-1 in control (Ctrl) and *spd-3(oj35)* (*spd-3*) exposed to *ima-2* (*RNAi)*. Times are in seconds relative to anaphase onset (t= 0). Scale bars, 10 μm. (**B**) Quantitation of normalized PLK-1::GFP intensity in the nucleoplasm of embryos from time-lapse sequences as in *(A)*. The value of PLK-1 intensity in control (red, n=3 for each condition) and *spd-3(oj35)* (purple, n=3 for each condition) without (solid) and with (empty) *ima-2 (RNAi)*. Error bars are SD. (**C-E**) Graph plotting percentage of SDC (**C**) and spindle misalignment (**D, E**) in control and *spd-3(oj35). ima-2 RNAi* is in red (n=3 for each strain), and *plk-1 RNAi* (n=6 for each strain) is in yellow.

**Figure S8.**
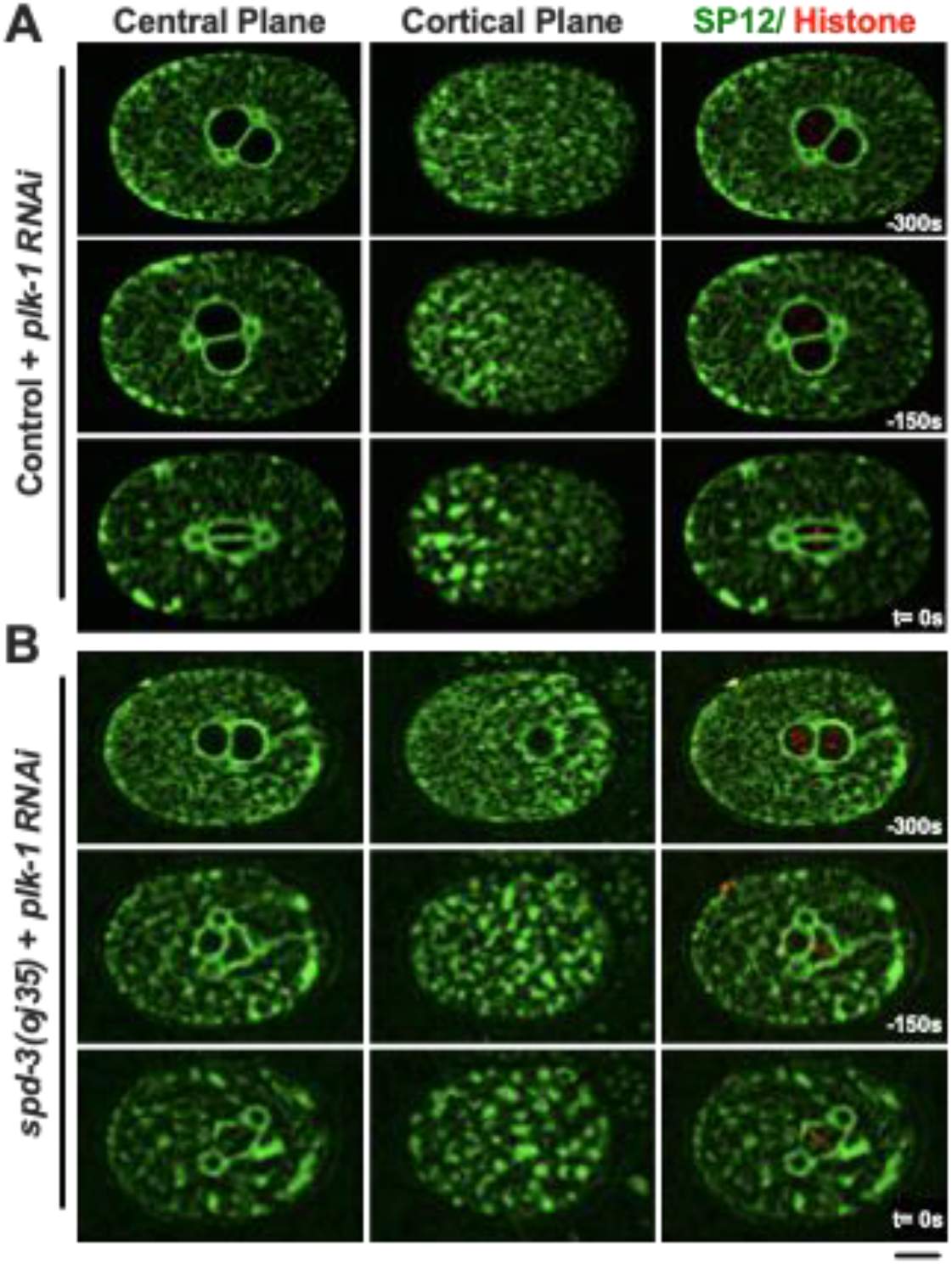
Changes in ER morphology are not induced by elevated PLK-1 levels, related to Figure 5. (**A**) Representative images of control embryos treated with *plk-1* (*RNAi*) coexpressing SP12::GFP and mCherry::histone H2B. Images of the ER marker SP12 acquired at the central (**left**) and cortical plane (**middle**) and a merge of SP12 and histone H2B (**right**) are shown. (**B**) The same layout of images as in (**A**) but showing the morphology of ER after *plk-1* (RNAi) in the *spd-3(oj35)*. Times are in seconds relative to anaphase onset (t= 0). Scale bars, 10 μm.

**Figure S9.**
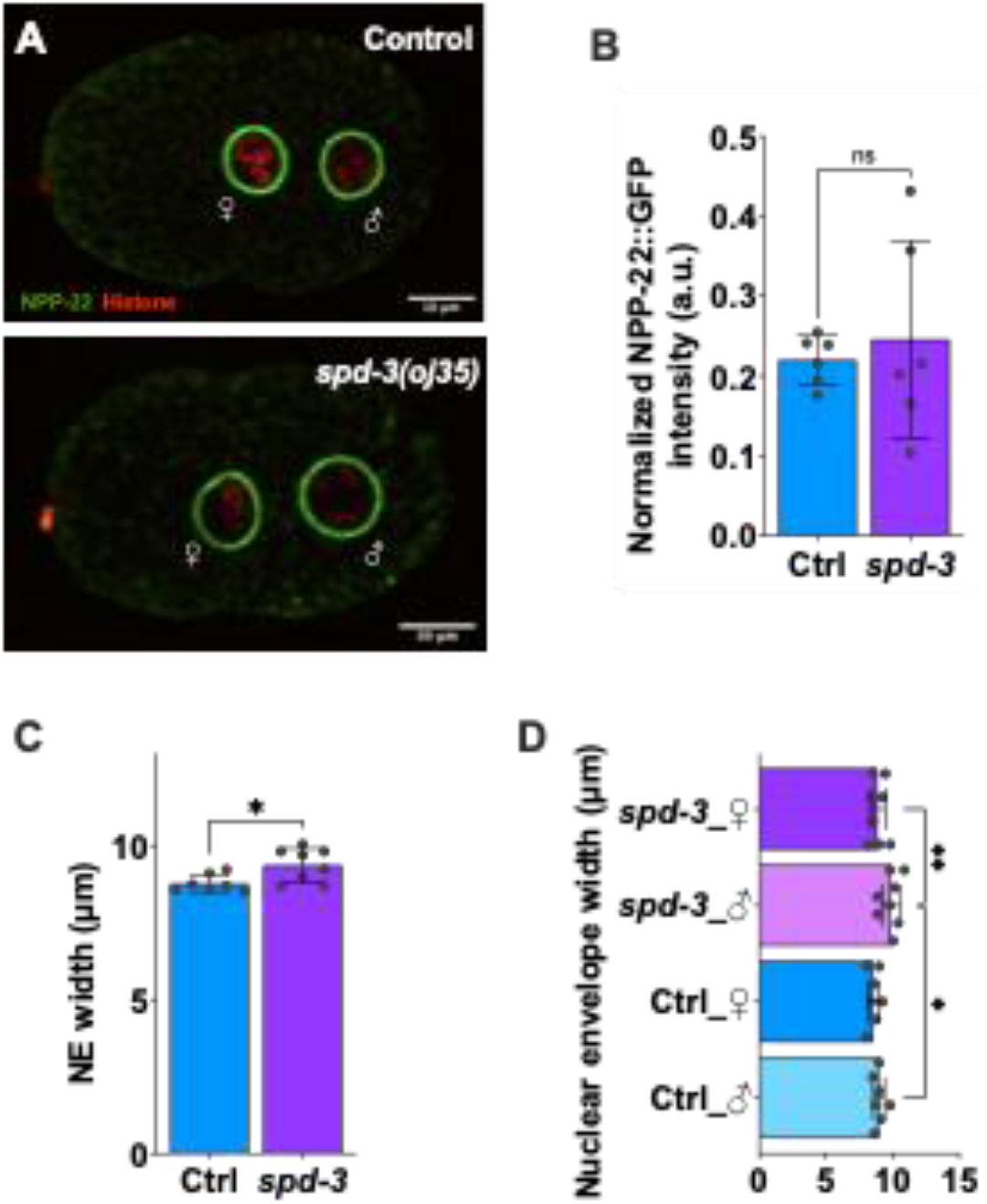
The male pronucleus expands before the pronuclear meeting, related to Figure 5. (**A**) Representative images of embryos coexpressing NPP-22::mNG and mCherry::histone H2B in control (Ctrl) and *spd-3(oj35)* (*spd-3*). Scale bars, 10 μm. (**B**) Plot the intensity of NPP-22::mNG in the nuclear membrane of control (n=7) and *spd-3(oj35)* (n=8). (**C**) Plot of the pronuclear envelope width in control and *spd-3(oj35)*. (**D**) Plot the nuclear envelope width of male and female pronuclei in control and *spd-3(oj35)*. (**B-D**) Error bars are SD. The significance of the difference between strains was determined by t-tests. ns indicates no significant difference. \**P*<0.05. \*\**P*<0.01.

**Figure S10.**
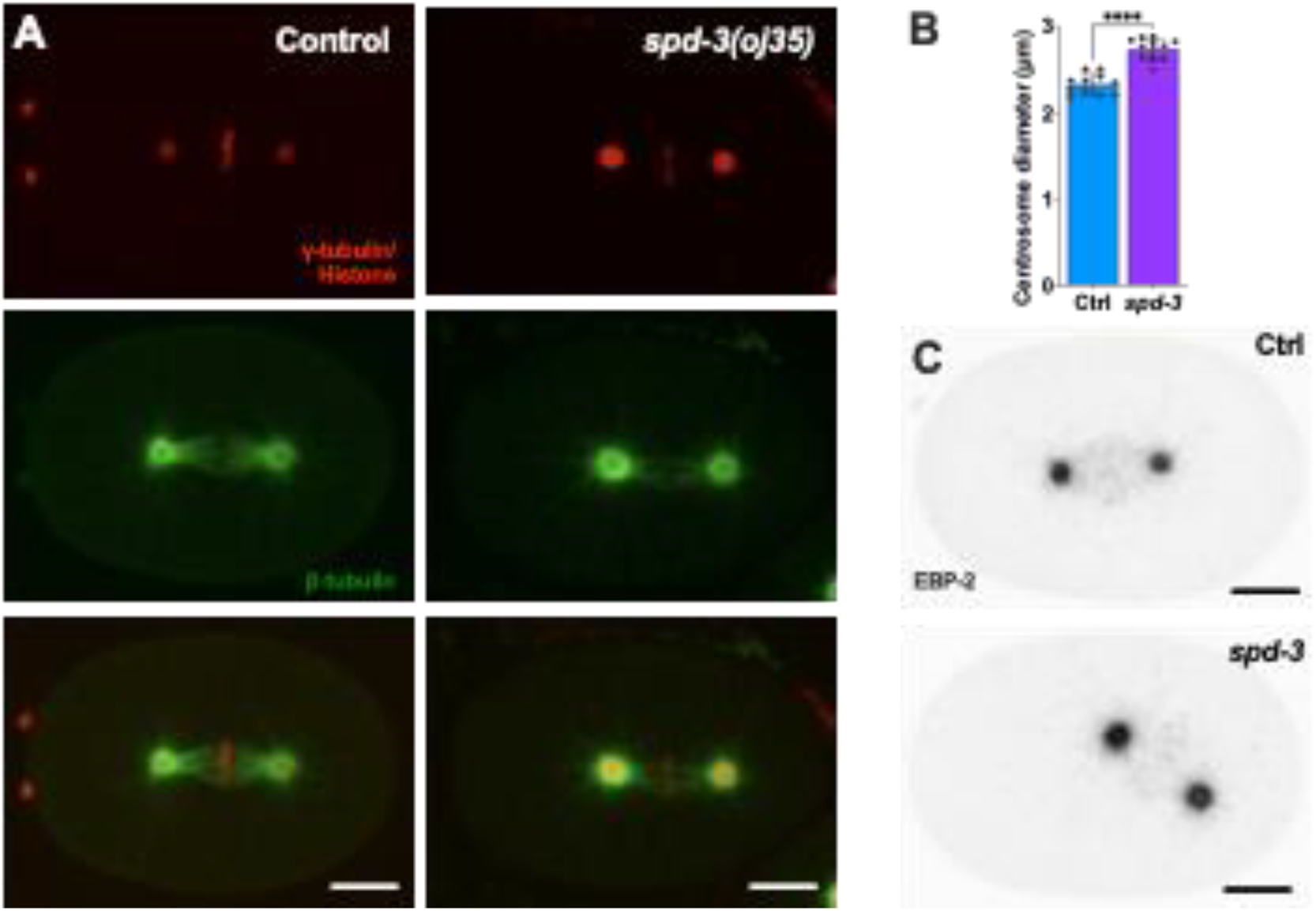
The pericentriolar material increases in size in *spd-3(oj35)*, related to Figure 5. (A) Spinning-disk confocal images of embryos in control (**left**, Ctrl) and *spd-3(oj35)* (**right**, *spd-3*). Embryos are coexpressing mCherry::γ-tubulin, mCherry::histone H2B, and GFP::β-tubulin. (**B**) Plots of the centrosome size in control (n=11) and *spd-3(oj35)* (n=11). The diameter of centrosomes was measured by the intensity of mCherry::γ-tubulin. (**C**) Spinning-disk confocal images of embryos in control (**left**, Ctrl) and *spd-3(oj35)* (**right**, *spd-3*). Embryos are coexpressing mCherry::histone H2B, and EBP-2::GFP. (**A, C**) Scale bars, 10 μm.

## SUPPLEMENTAL MATERIALS AND METHODS

### *C. elegans* strains used in this study

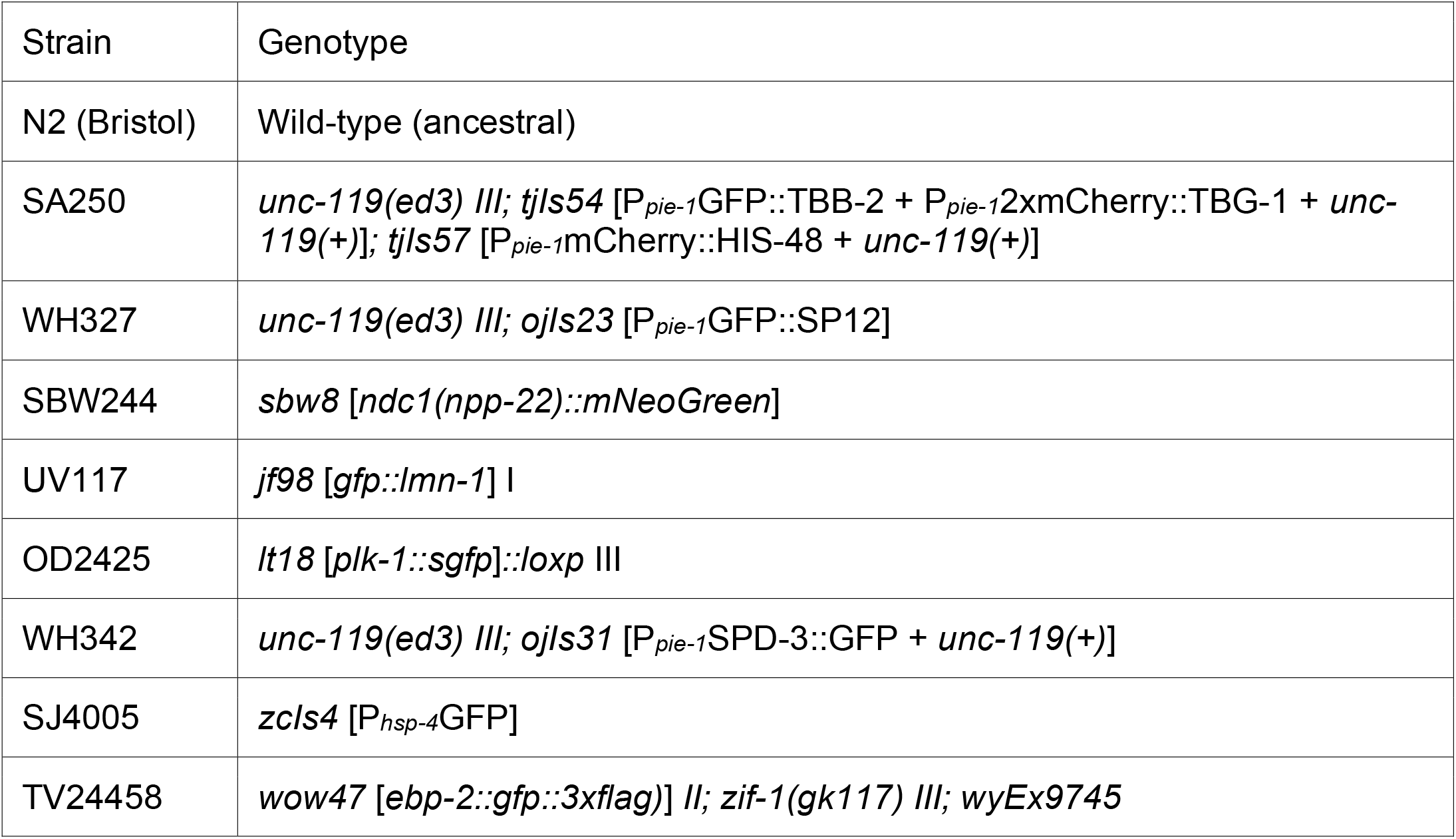

The *C. elegans* strains used in this study are listed in the table above. The *spd-3(oj35)* allele was outcrossed against N2 five times before crossing into fluorescent marker strains for phenotypic analysis.

### *C. elegans* strains and alleles

The Bristol strain N2 was used as the standard wild-type strain. Culturing, handling, and genetic manipulation of *C. elegans* were performed using standard procedures (Brenner 1974). All *C. elegans* strains and the temperature-sensitive strain *spd-3(oj35)* were maintained at 16°C and L4 hermaphrodites were shifted to room temperature (~23°C) overnight and heat-shocked at 25°C for 4 hr prior to analysis. The following alleles and strains were used: LGIII: *clk-1(qm30);* LGIV: *isp-1(qm150), spd-3(oj35)*. They were backcrossed four times with wild-type animals before being used for genetic analyses.

#### *C. elegans* RNA-mediated interference

Primers for *C. elegans* RNAi feeding clones

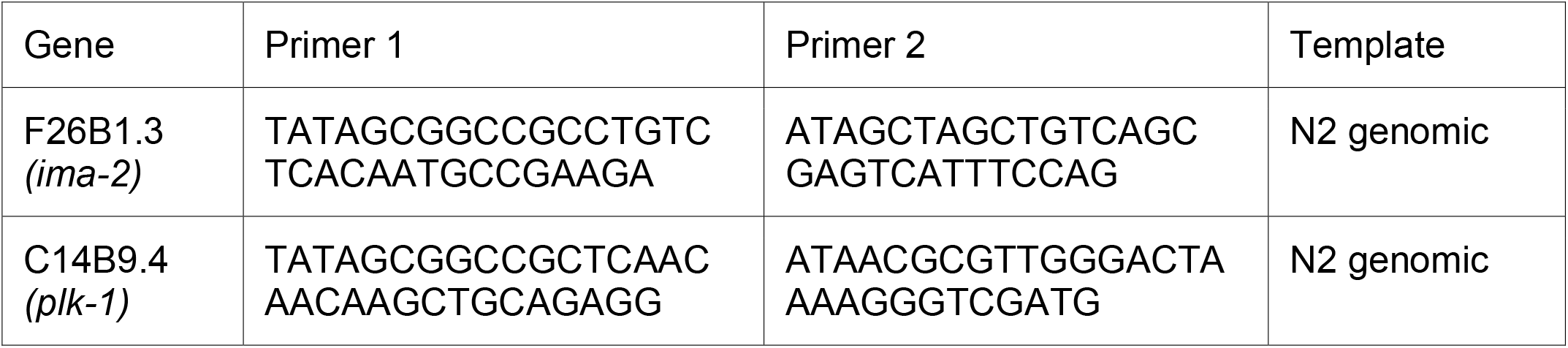

### RNA-mediated interference

The *ucr-1, mev-1, cco-1, mtx-1, gop-3, yop-1, hsp-3, hsp-4, ima-2* and *plk-1* (RNAi) were administered by feeding. Feeding vector (L4440) was obtained from the Addgene. The *ima-2 and plk-1* (RNAi) feeding vectors were constructed by cloning the *ima-2* and *plk-1* genomic DNA into the L4440 feeding vector, followed by transformation into *Escherichia coli HT115(DE3)* bacteria. The *E. coli HT115(DE3)* bacteria containing the feeding vectors were cultured and used to seed RNAi plates (1 mM IPTG and 50 μg/ml ampicillin). L4 hermaphrodites were allowed to feed for 24, or 48 hr at room temperature (~23°C) and then shift to 25°C for 4 hr before analysis. For *plk-1* RNAi, L4 larvae were cultured at room temperature overnight and then transferred to *plk-1* RNAi feeding plates allowed to feed for 6 hr at 25°C before analysis.

### Live imaging

Embryos for live-imaging experiments were obtained by dissecting gravid adult hermaphrodites in M9 buffer (42 mM Na_2_HPO4, 22 mM KH_2_PO_4_, 86 mM NaCl, and 1 mM MgSO_4_). One-cell embryos were mounted on slides with 2% agarose pad, overlaid with a 22 × 22-mm coverslip, and imaged at room temperature on fluorescence microscope (ECLIPSE Ti2; Nikon) equipped with a CFI Apo TIRF 60× 1.45 NA oil immersion lens and a CCD camera (iXon 897 Ultra EMCCDs) or on a 3i VIVO spinning-disc confocal microscope (Axio Examiner.Z1; Zeiss) equipped with Zeiss Plan-Apochromat 63x/1.40 oil microscope objective, 6 laster lines and a Hamamatsu ORCA-Flash4.0 scientific CMOS camera (Hamamatsu) for detection. For z-stacks, images in a 27-micron z-series (1 micron per section) were captured every 15 s (Figure 1A, B, D, E; Table S1; Fig. S2; Fig. S5; Fig. S10A). The microscope was controlled by Nikon NIS-Elements software (Nikon). Fluorescence confocal images were acquired every 10 s (Figure 4, 5; Fig. S8; Fig. S9) or 15 s (Figure 1-3; Fig. S4; Fig. S6; Fig. S7) by collecting 31 z-planes at 1.0-µm intervals or 17 z-planes at 1-µm intervals without binning. Imaging was initiated in one-cell embryos before pronuclear meeting and was terminated 3 min after anaphase onset. EBP-2::GFP images acquired at 400 msec intervals and each movie was 150 frames per 1 min (Fig. S10C). Acquisition parameters were controlled using a Slidebook 6.0 program (3i - Intelligent Imaging).

### Image analysis

All images were processed and analyzed using ImageJ (National Institutes of Health). For figure construction, final image panels were scaled for presentation in Prism v9.5.0 (GraphPad). Lamin fluorescence intensity (GFP::LMN-1) was quantified using ImageJ software by drawing a box around the nuclei and subtracting the background fluorescence in an equal-sized box drawn over the cytoplasm (Fig 3C and D). To quantify the PLK-1::GFP fluorescence in nucleoplasm, a fixed-size box was drawn in nucleoplasm (red box) and in cytoplasm (black box) as background at each time point. The normalized PLK-1 level was calculated as [(integrated intensity in red box – integrated intensity in black box)/ integrated intensity in black box] (Fig. 4C and D; Fig. S7B). A gamma of 1.5 was applied for images of centrosomal PLK-1::GFP (Fig. 4E) to visualize centrioles in the mutants, while not oversaturating the signal in the WT centrosomes. The fluorescence of *hsp-4::ghp* reporter was measured using ImageJ software by drawing a box around the individual worm for quantification and subtracting the background fluorescence in an equal-sized box drawn over the background fluorescence (Fig. S5). For centrosome size, centrosome was marked with mCherry::γ-tubulin and line scans from single confocal planes over centrosome regions, allowing quantification of the centrosomes diameter and the full width at half maximum taken as centrosome diameter (Fig. S10).

### Fluorescence lifetime imaging (FLIM) microscopy

A Zeiss LSM-780 NLO confocal/multiphoton microscopy system consisting of an inverted Axio Observer (Zeiss) microscope, motorized stage for automated scanning, Chameleon Vision-II (Coherent Inc.) ultrafast Ti:sapphire laser with dispersion compensation to maintain pulses at the specimen plane (690– 1,060 nm, 80 MHz, 150 fs) for multiphoton excitation and a standard set of dry and immersion objectives was used. Three HPM-100-40 hybrid GaAsP detectors (Becker and Hickl) are connected to the nondescanned (NDD) port of the microscope using two T-adapters (Zeiss) with proper dichroics and band pass filters to collect as much fluorescence as possible in the spectral ranges tryptophan channel: 740 nm Ex, 340–380 nm Em, NAD(P)H channel: 740 nm Ex, 460–500 nm Em and FAD channel: 890 nm Ex, 520–560 nm Em. The three channels also contain a 690 nm short pass filter (Zeiss) in the beam path to avoid excitation background. Three SPC-150 cards (Becker and Hickl) synchronized with the pulsed laser and the Zeiss LSM-780 scan head signals collect the time-resolved fluorescence in TCSPC mode using SPCM (Version 9.74) acquisition software. A motorized stage is used during imaging using Zeiss 40x NA1.3 oil apochromatic objective lens.

#### FLIM Processing and Analysis

FLIM data fitting is based on the B&H handbook. We used a two components incomplete model to fit Tryptophan, NAD(P)H and FAD/FMN FLIM images. The offset and scattering are set to 0. Shift is optimized to make sure the Chi2 as close as to 1.

The FLIM processing followed the previously published paper (Wallrabe et al. 2018; Cao et al. 2019) with an important advance in normalization of photon reference images to compensate for varying intensities (FIJI, Plugins -> Integral Image Filters -> Normalize Local Contrast): followed by zero’ing the nucleus; cell segmentation and creating single pixel ROIs by a ImageJ/FIJI custom plugin. The purpose of this sequence is to create pixel locations by X-Y coordinates, specific for embryos. Those locations are then applied to the FLIM data to extract any of the FLIM parameters in the data pool. A custom Python code ultimately analyzes different data combinations to produce ratios, means, medians, and histograms, further charted in MS Excel.

### Immunoblotting

200 adult worms from each strain were transferred to 30 μl of sample buffer (80.0 mM Tris pH 6.8, 2.0% SDS, 10.0% glycerol, 0.0006% Bromophenol blue, and 5.0% β-mercaptoethanol), subjected to the freeze-thaw treatment three times in liquid nitrogen and water bath and then boiled in a 95°C-heating block for 5 minutes. The samples were loaded on 10% SDS-PAGE, transferred to a polyvinylidene fluoride (PVDF) membrane and subjected to western blot analysis using anti-PLK-1 (1:3,000 dilution, from Monica Gotta lab). Primary antibodies were detected using IRDye® 680RD goat anti-Rabbit IgG secondary antibody (1:5,000 dilution; Abcam) and then probed for α-tubulin using the monoclonal DM1α antibody (1:3,000 dilution; Sigma Aldrich) to examine the expression levels of α-tubulin as a loading control. PLK-1 western blots were quantified with ImageJ (Fig. 4B and Fig. 5B). Antibody signals were detected using the Azure Biosystems c600 and Li-COR Fc Odyssey imaging system.

### Statistical analysis

Statistical analysis was conducted using Prism v9.5.0 (GraphPad). P values were determined using unpaired two-tailed t tests assuming equal SD. P > 0.05 (ns), P < 0.05 (*), P < 0.01 (**), P < 0.001 (***), and P < 0.0001 (****). Data distribution was assumed to be normal, but this was not formally tested.

